# Ecological bleaching trajectories under severe heat stress are only partially captured by acute heat stress assays

**DOI:** 10.64898/2026.05.14.725291

**Authors:** Sebastian Szereday, Kok Lynn Chew, Joseph A. Henry, Natasha Zulaikha, Christian R. Voolstra

## Abstract

Global marine heatwaves have devastated tropical coral reefs, and further mortality is projected under ongoing climate change. Identifying thermally tolerant coral colonies is therefore a priority for conservation, restoration, and research. Portable acute heat stress assays (e.g., CBASS) enable rapid, standardized estimates of coral thermal tolerance under field conditions. However, it remains unresolved whether such experimentally derived metrics (ED5, ED50, DW) predict bleaching and mortality in situ. Here, we quantified acute thermal tolerance metrics for 2,068 coral colonies across 12 common Indo-Pacific species, six months prior to an unprecedented heat stress event in northeastern Peninsular Malaysia and compared experimentally derived ED and DW values to subsequent bleaching severity and mortality in the field. Experimental thermal tolerance metrics explained only a limited proportion of variation in bleaching outcomes and survival. Predictive power varied among species and was higher in slow-growing species. Our findings suggest that while acute heat stress assays capture substantial variation in coral thermal tolerance, their ability to predict in situ outcomes is context-dependent and diminishes under severe thermal stress. Ultimately, in situ coral bleaching under severe heat stress may reduce the discriminatory capacity of acute assay-derived tolerance metrics.

## Introduction

Climate change has significantly warmed the world’s oceans, resulting in more frequent, longer, and more widespread marine heatwaves^1,2^. This exposes marine ecosystems to unprecedented heat stress with detrimental effects on marine life and coastal communities. On tropical coral reefs, heat stress disrupts the coral–algal symbiosis, leading to bleaching and increased susceptibility of reefs to climate-driven mortality^3,4^. The world’s coral reefs have now experienced four global coral bleaching events in less than three decades^5^, resulting in severe loss of coral reef cover, abundance, and functioning^6,7^. Consequently, an estimated 50% of coral reef has been lost since the mid-20^th^ century, threatening the ecological and economic services that coral reefs provide^8^. In line with this, the most recent report from the Stockholm Resilience Centre declared that coral reefs have passed a tipping point, which must be understood as an urgent call to action. Given that current global policy seems insufficient to prevent further mass coral mortality under ongoing climate change, a key challenge is to identify and select thermally tolerant coral genotypes for conservation and restoration efforts aimed at enhancing reef resilience^9^. These efforts may include assisted evolution^10,11^, stress hardening^12^, and microbiome-based interventions^13^, ultimately aiming at the functional rebuilding of coral reefs^14^. Identifying thermally resilient corals, however, remains a logistical and biological challenge^15,16^, creating a need to develop technologies capable of standardized, high throughput, and easy to deploy experimental systems to efficiently and robustly identify heat tolerant phenotypes in the field across remote locations.

Identifying heat tolerant corals can be achieved by tracking colonies in situ during and across bleaching events. Such temporal surveys are logistically demanding and rely on in situ marine heatwaves that induce ecologically relevant bleaching and mortality to allow differentiating resilient colonies, more likely to survive future bleaching events. Importantly, evidence suggests that bleaching susceptibility varies across time and space^16–18^, which may obscure accurate selection of heat tolerant individuals. Further, bleaching susceptibility per se is not an indicator of thermal tolerance^19^, as resilience can manifest in the form of bleaching and subsequent recovery vs. the inability of a coral to bleach until mortality is reached^20,21^. To this end, ex situ heat stress experiments have been used to screen corals to identify heat tolerant populations, species, and superior host-algal combinations^22,23^. Because coral bleaching involves temperature-dependent impairment of photosystem II (PSII) in algal symbionts^24,25^, retention of maximum photosynthetic efficiency (i.e., F_v_/F_m_) under heat stress has been widely used as a proxy for bleaching susceptibility^26^. Accordingly, changes in F_v_/F_m_ across different temperature stress exposures provide a representative and non-invasive measure of coral thermal tolerance for experimental coral biology^27,28^. Following this, acute heat stress assays rapidly quantify F_v_/F_m_ in the field, showing promising results in identifying thermally tolerant corals for in situ reef restoration^29,30^. However, a standardized methodology and operational system were missing until recently^31^, preventing cross-study comparison across regions, reefs, species, and coral colonies.

The Coral Bleaching Automated Stress System (CBASS) was conceived as a low-cost portable system developed to yield standardized coral thermal tolerance metrics within hours in remote filed location^22^. Due to its standardized operating platform and methodology, it has become an established and widely used screening tool to investigate coral thermal tolerance^32–34^. Initial CBASS studies demonstrated the ability to resolve ecologically observed coral bleaching differences across reef sites^27^, regions^35^, and populations^36^ and to capture thermal tolerance phenotypes that are broadly consistent with patterns observed in longer-term experiments^27^. These proof-of-principle studies highlighted the potential of rapid phenotypic diagnostics to identify coral colonies with elevated thermal tolerance, thereby contributing to the proliferation of acute heat stress assay studies. For instance, thermal tolerance metrics determined from acute heat stress assays were correlated with symbiotic, molecular, and environmental factors to elucidate the underpinnings of thermal resilience^21,37–42^. Importantly, for coral restoration, CBASS assays were used to delineate genotype-level heat tolerance differences within coral nursery populations^43^, and to support the selection of heat tolerant corals for reef restoration^29,44^.

Despite the widespread application of acute heat stress assays, it remains unresolved to what extent experimentally derived thermal tolerance metrics—such as the onset of stress (ED5), the temperature tolerance threshold (ED50), and the upper thermal limit (ED95) — predict bleaching outcomes in situ^23,29,33,45^. This knowledge gap requires urgent investigation across reef scales and species with differential morphology and life-history strategies to determine whether corals exhibiting superior heat stress tolerance in acute heat stress assays also exhibit reduced severe bleaching and mortality under ecologically relevant conditions. Here, we analyse data from over 1,850 coral colonies to evaluate how well experimentally derived thermal tolerance metrics predict in situ bleaching outcomes across 12 Indo-Pacific reef-building species. Thermal tolerance metrics were quantified approximately six months before the most severe heat stress event recorded in the study region. Our results show that experimentally derived thermal tolerance metrics do not consistently predict in situ bleaching outcomes, with predictive strength varying among species. On the one hand, this highlights that bleaching susceptibility cannot be equated with experimentally determined thermal tolerances, reflecting the ecological complexity of bleaching responses. On the other hand, it cautions that under sufficiently high thermal stress, bleaching responses converge, diminishing the influence of stress tolerance differences, holding important implications for coral conservation and restoration strategies.

## Methods

### Research site and coral colony sampling

The research was conducted between September 2023 and November 2024 in Pulau Lang Tengah (5.793844, 102.895492), in northeastern Peninsular Malaysia (Supplementary Figure 1), a biodiverse reef region of ecological and economic significance where coral research and restoration efforts are ongoing^46,47^. To account for reef variability and to obtain a population level representation, 2,068 coral colonies of 12 species with varying morphologies and physiologies were randomly selected from five reef sites with distinct micro-environments (i.e., differential exposure to winds and waves, varying diel temperature regimes, and differences in benthic substrate composition and reef complexity, sensu Bernard et al. 2023^48^ and Szereday et al., 2024^49^). Heat stress assays were carried out daily between September 11, 2023, and November 4, 2023 (further described below). For this purpose, each morning before acute heat stress assays, four replicate fragments of approximately 6.0-9.0 cm^2^ size were collected from each coral colony from four opposing sides with garden shears for acute heat stress testing. For massive coral species, 40 mm diameter cores were removed using an underwater power drill fitted with diamond hole-saw drill bit. For each colony, fragments were placed in a numbered zip-lock bag (for identity tracking) and transported in a bucket with local sea water to the ex situ experimental CBASS (approximately 10 minutes from the furthest reef site by speed boat). Concurrently, upon underwater sampling, coral colonies were tagged with numbered rubber tags, photographed, and mapped to ensure re-detection and identification during subsequent coral bleaching surveys between March and October 2024. All colonies were visually healthy at the time of sampling, i.e., showed no signs of disease, bleaching, partial mortality, or predation. Further, at each reef site, selected colonies were proportionately selected from a broad depth range between 4-16 m to maximize phenotype diversity (except for *Acropora* cf. *spicifera* whose depth range is 2-6 m, ±1 m tidal range), and colonies were spaced at a minimum distance of 5 m to avoid sampling of clonal genotypes^50^.

### Standardized acute heat stress assays (CBASS)

To derive standardized experimental heat stress metrics (ED5: thermal onset; ED50: thermal tolerance threshold; ED95: thermal limit) in the field, we employed the Coral Bleaching Automated Stress Systems (CBASS)^21,22,27,32^ for rapid and high-throughput acute heat stress testing of selected coral colonies. Each day between September 11, 2023, and November 4, 2023, two to three independent CBASS systems were operated in parallel with an average of 16 different colonies from the same species per CBASS run. A total of 126 CBASS assays were run to test the thermal tolerance of 2,068 coral colonies. Each CBASS system consisted of four replicate 10 L flow-through tanks with independent temperature profiles customized to prevailing local reef conditions. The selected temperature profiles were based on available NOAA climatology (Coral Reef Watch, version 3.1., 2014), binned to the nearest 5 km satellite pixel (Liu et al., 2014). The control baseline temperature was set at 30°C, based on NOAA’s historical maximum monthly mean (MMM) temperature of 29.94 °C. The three heat-ramp-hold temperature profiles were then set to MMM+4°C (34 °C), MMM+8°C (38 °C), and MMM+12°C (42 °C) to induce mild, strong, and severe heat stress, respectively. Of note, these temperature profiles were selected based on previous CBASS assays conducted at this location^29^, as well as based on a brief pilot study in September 2023 to determine the heat-hold temperatures that induce sufficient heat stress to satisfy the assumptions of subsequent dose-response curve (DRC) modelling, i.e., increasing photosynthetic efficiency decline (further details below) with increasing temperature^23^.

The four replicate fragments from each coral colony were distributed across the four temperature treatment tanks, so that each coral colony was subjected to each of the four temperature profiles. Fragments were fixed on aquarium-trays corresponding to their in situ orientation (i.e., either fixed upright or laid down horizontally). Following the placement of all fragments, each tray with coral fragments was photographed with a Coral Watch Coral Health Chart^51^. The 7h thermal cycles started at 1 pm and consisted of a 3h heating phase to the desired heat stress temperature, a 3h heat-hold phase of the maximum treatment temperature, and a 1h cooling phase back to the baseline temperature (Supplementary Figure 2). Temperatures in each treatment tank were controlled with temperature controllers (InkBird ITC-310T-B) that regulated thermo-electric chillers (IceProbe, NovaTec) and 200W titanium heaters (Schego). Light was supplied with dimmable 165W full spectrum LED aquarium lights, set at a 50-50 ratio of blue and white light, respectively, to match in situ reef conditions of approximately 35,000 lux intensity (i.e., ∼ 540–580 µmol photons m⁻² s⁻¹). Temperature in each treatment tank was measured every minute with HOBO MX2202 Pendant loggers (Onset Computer Corporation, USA, accuracy level ±0.5°C). Each morning, local seawater was collected and adjusted to baseline temperature (i.e., 30 °C) in three 200 L insulated reservoir tanks (one for each CBASS system) before the start of the CBASS assays. Once the desired temperature was reached, all treatment tanks were filled, and seawater was continuously supplied to treatment tanks with a submersible aquarium pump at a rate of 25-30 mL/min to achieve 100% turnover of each temperature tank volume every six hours. During the 1-hour cooling phase (starting daily at dusk at 6 pm), the white LEDs were first switched off, then the blue LEDs (30 min later), so that all fragments were dark-acclimated, completing the 7h diel temperature cycle. At this point, dark-acclimated maximum photosynthetic efficiency of PSII (F_v_/F_m_) of the coral algae was measured by pulse amplitude modulated (PAM) fluorometry (Diving-PAM underwater fluorometer; Heinz Walz GmbH, Effeltrich, Germany) to derive a physiological measure reflective of heat stress tolerance of corals^26,28^. Two F_v_/F_m_ measurements were recorded per fragment from two separate spots of the fragment surface, and a mean F_v_/F_m_ value per fragment per temperature treatment was calculated. To cross-check consistency, the standard deviation of F_v_/F_m_ measurements for each temperature treatment, species, and site combination was calculated (i.e., Species 1, Site 1, Treatment 30 °C, Species 1, Site 1, Treatment 34 °C, etc.). Then, for each fragment, the difference between the two measured F_v_/F_m_ values was calculated (i.e., value 1 - value 2), and the higher value was removed if the difference was >1 standard deviation from the mean of all measurements. Finally, a coral fragment was considered dead if three consecutive F_v_/F_m_ measurements failed (i.e., no photosynthetic signal or F_v_/F_m_= 0.00). Following all F_v_/F_m_ measurements, aquarium trays were photographed again using the same camera with a Coral Watch Coral Health Chart as a colour reference standard.

### Standardized experimental heat stress metrics

Standardized thermal tolerance metrics were calculated based on photochemical yield measurements (F_v_/F_m_) obtained during CBASS assays. Effective dose 50 (ED50) values represent standardized thermal tolerance thresholds^23^. ED50 is defined as the temperature at which the measured parameter (here F_v_/F_m_), decreased by 50% of the initial value at baseline temperature, obtained through log-logistic regression curves from measured parameter declines over temperature increases. Additional ED values can be obtained from the curve fit for a more detailed assessment of the coral heat stress response^28^. Therefore, we also determined the effective dose 5 (ED5) value, which signifies the thermal breakpoint temperature, i.e., the onset temperature of thermal stress at which a decline in photosynthetic efficiency is initiated. Further to this, the ED95 value, which signifies the thermal limit, i.e., the temperature at which 95% of photosynthetic efficiency is lost, was obtained to calculate the Decline Width (DW) of the fitted F_v_/F_m_ curve, which is calculated as ED95-ED5^42^. The Decline Width describes whether the effective loss of photosynthetic efficiency occurs gradually or rapidly (i.e., higher values suggestive of higher tolerance). For each colony, photochemical yield measurements were fitted on log-logistic dose-response curves with the DRC package in R^52^ using the three-parameter model (LL.3) with model limits for the three parameters ‘hill’, ‘max’, and ‘ed50’ (upper limits=100, 0.8, NA; lower limits=10, 0.3, 30). Since DRC modelling assumes a steady decline in the measured parameter (i.e., F_v_/F_m_) with increasing temperature, i.e., across temperature treatments, we removed a large number of colonies from ED and DW modelling, as they did not conform to the expected pattern (see Table 1). We excluded any coral colony from the dataset whose F_v_/F_m_ value did not decrease with increasing temperature, but we allowed for minor variation (±0.05) such that small increases or stable values between treatments were treated as effectively unchanged. To utilize the entirety of the dataset outside the dose-response curve modelling constraints, we also calculated the relative change in F_v_/F_m_ measurements between the baseline and the two highest temperature treatments (i.e., ΔF_v_/F_m_ 38°C and ΔF_v_/F_m_ 42°C). Explicitly, we calculated the decline in F_v_/F_m_ values between temperature treatments by calculating ΔF_v_/F_m_ 38°C = [(F_v_/F_m_ 30°C - F_v_/F_m_ 38°C) / F_v_/F_m_ 30°C] and ΔF_v_/F_m_ 42°C = [(F_v_/F_m_ 30°C - F_v_/F_m_ 42°C) / F_v_/F_m_ 30°C].

**Table 1.**
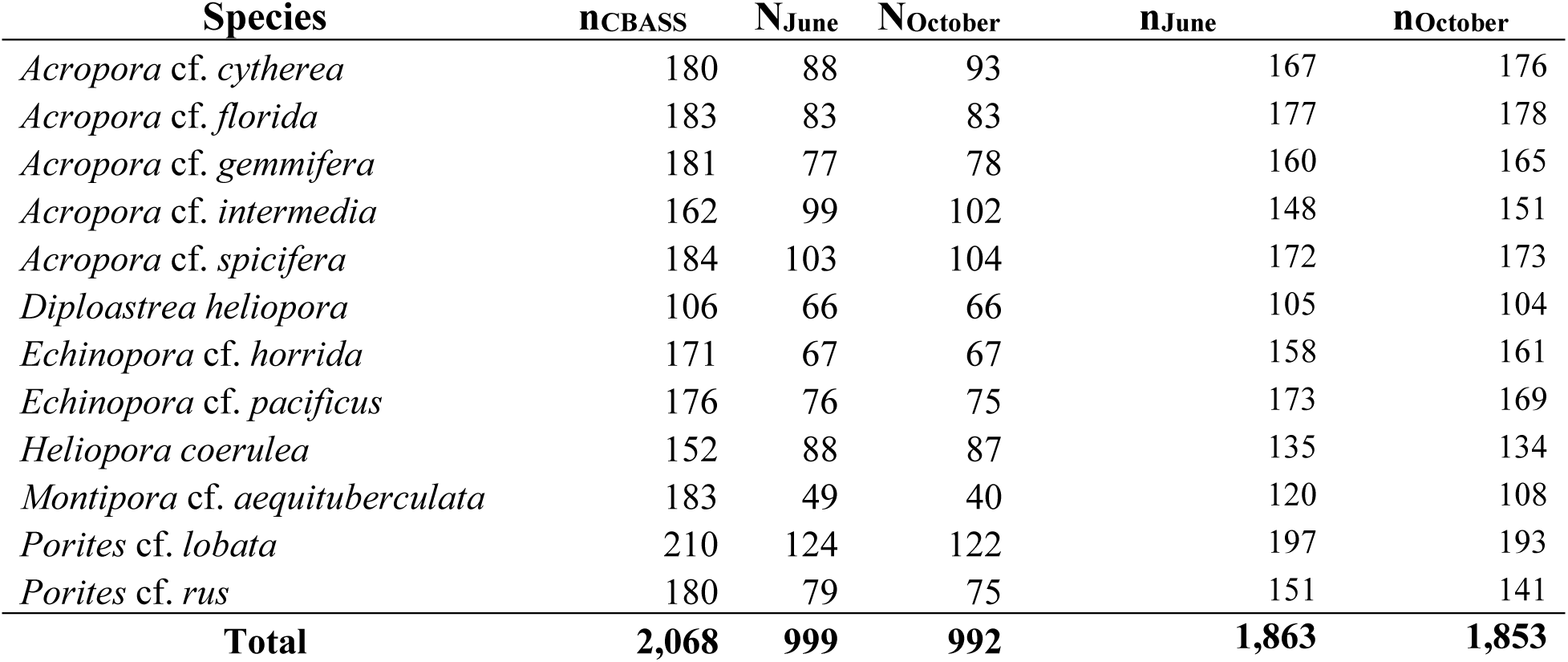
Bleaching and experimental heat stress dataset. Experimental heat stress metrics (i.e., n_CBASS_) were assessed for 2,068 colonies between September and November 2023. Of these, a total of 1,863 (i.e., n_June_) and 1,853 (n_October_) colonies were surveyed for *in situ* bleaching in June 2024 and subsequent post-bleaching mortality in October 2024, respectively. The differences in colony numbers between n_June_, n_October_, and n_CBASS_ are due to non-detection of colonies during in situ bleaching surveys (i.e., colony not found). N_June_ and N_October_ are the number of coral colonies with determined standardized heat stress metrics based on photochemical efficiency (i.e., F_v_/F_m_) and their corresponding *in situ* bleaching data. The difference in colony numbers between n_CBASS_ and N_June_/ N_October_ is due to the dose-response curve (DRC) model constraints, whereby photosynthetic efficiency (i.e., F_v_/F_m_) was bound to decline with increasing temperature for accurate modelling, with non-confirming colonies being excluded (∼ half of all colonies). Experimental heat stress metrics not based on DRC modelling (i.e., visual bleaching score and ΔF_v_/F_m_ changes between temperature treatments) were assessed for all colonies that were acute heat stress tested and surveyed for in situ bleaching (i.e., n_June_ and n_October_).

To complement pulse amplitude modulated (PAM) fluorometry-based thermal tolerance assessments, and because in situ bleaching assessments predominantly assess bleaching severity visually^53^, a visual Bleaching Index Score (BIS) was calculated following the methodology by Alderdice et al. (2022)^54^. Specifically, colour scores of the coral tissue of CBASS’ed fragments were assigned by two independent scorers based on reference to a colorimetric reference card (i.e., Coral Watch Health Chart) on a six-point scale, with a score of 6 representing maximum pigmentation and a score of 1 representing no pigmentation (i.e., complete bleaching). Colour scores were assigned for each fragment before and after the CBASS assays (i.e., 2 scores per fragment x 4 fragments per colony x 2,068 colonies), and the relative decline in pigmentation (i.e., colour score before – colour score after) was calculated as a measure of visual bleaching. Given the DRC constraints, as well as the coarse resolution of this approach (i.e., coral colour is a discrete variable with non-continuous values), we did not subject the scores to DRC modelling. Rather, the two values from the two independent assessors were averaged, and this value was used to calculate the ΔBIS between the control treatment (i.e., 30°C) and the two highest temperature treatments 38°C and 42°C, based on the same formula as above, i.e., ΔBIS 38 °C = [(BIS 30 °C - BIS 38 °C) / BIS 30 °C] and ΔBIS 42°C = [(BIS 30°C - BIS 42°C) / BIS 30°C].

### In situ heat stress and coral bleaching monitoring

Sea surface temperature (SST) data for Pulau Lang Tengah were sourced from the National Oceanic and Atmospheric Administration’s (NOAA) Coral Reef Watch (CRW) version 3.1, binned to the nearest 5 km^2^ satellite pixel (5.793844, 102.895492)^55^. The common heat stress metric, Degree Heating Weeks (DHW, described as °C-weeks)^56^ was used as a measure of heat stress accumulation. Additionally, we used a more sensitive (sensu DeCarlo, 2019^57^) DHW metric, which better reflects local bleaching thresholds (i.e., nDHW, see Szereday et al., 2024^49^) to describe heat stress levels between January 1, 2023, and November 1, 2024. To measure reef-scale in situ heat stress across depths, ten HOBO ProV2 temperature loggers (Onset Computer Corporation, USA, accuracy level ±0.2°C) were deployed at five sites at 5 meters and 15 meters in March 2024 (one per site per depth), logging at 60 minutes intervals. Following interpolation of the satellite-based MMM with in situ data based on protocols described by Szereday et al. (2025)^19^, nDHW was calculated based on in situ temperature data to determine heat stress levels across depths.

All study reef sites and tagged colonies were monitored for coral bleaching between March and November 2024. The bleaching analysis focused on survey timepoints in June 2024, reflecting the immediate heat stress response during peak heat stress, and in October 2024, reflecting the short-term recovery and mortality, noting that the October timepoint is of greater importance to identify ecologically relevant bleaching outcomes (e.g., survival vs. mortality). The severity of bleaching was assessed visually underwater on a per-colony level, noting the various bleaching severity levels (i.e., healthy, pale, bleached, dead, etc.) in assessment steps of 5%, thereby estimating the proportion of coral tissue bleached, whereby the sum of all observations added up to 100% (i.e., the entirety of the colony). In this way, within-colony bleaching heterogeneity, i.e., partial bleaching and mortality^58^, was accounted for. The severity of bleaching was then assessed by calculating the Colony Bleaching Response Index (CBRI)^19^, a normalized and weighted measure on a 0-100 scale, whereby higher values indicate more severe bleaching and bleaching induced mortality weighs the highest (i.e., each mortality increment of 5% increase the CBRI by 5). Of note, June CBRI values indicate bleaching severity during peak heat stress, and October CBRI values captured post-bleaching mortality (as further shown below). All colonies were photographed for reference during surveys using Olympus TG6 and TG7 underwater cameras.

### Data analysis

All statistical analyses were conducted at significance levels of p ≤ 0.05 at the species level for both time points (i.e., CBRI in June and CBRI in October). First, to resolve the resolution required to distinguish bleaching tolerant and susceptible colonies based on experimental thermal tolerance metrics ED5, ED50, and DW, pairwise Dunn’s Test with Benjamin-Hochberg corrections was used to calculate significant differences between colonies classified into quantile groups (i.e., top 20^th^ percentile, top 40^th^ percentile, top 60^th^ percentile, bottom 40^th^ percentile, and bottom 20^th^ percentile). Secondly, to explore whether colonies with higher thermal stress tolerance metrics also exhibited less in situ bleaching, we used a general linear model (GLM)^59^ to estimate the maximum likelihood of in situ bleaching at colony level as a function of ED50, ED5, and DW values, while considering ‘Depth’ and ‘Site’ as environmental covariates. For the purpose of modelling, coral colonies were ranked into quantiles based on their ED5, ED50, and DW values, respectively (i.e., top 20^th^ percentile, top 40^th^ percentile, top 60^th^ percentile, bottom 40^th^ percentile, and bottom 20^th^ percentile), and a logit-link function was incorporated into the model to transform the linear predictor into a probability scale between 0–1. All GLM analyses were performed in R Studio using the packages ‘*modEVA*’ and ‘*car*’. Estimated probabilities were extracted from the model to visualize how predicted bleaching (i.e., June data point) and post-bleaching mortality (i.e., October data point) differ across quantile groups. To further corroborate the modelled pattern with observed bleaching severity and post-bleaching mortality in the field, mean differences between groups were tested using pairwise Dunn’s test with a Benjamin-Hochberg correction. Hereby, the mean CBRI for June and October was calculated for colonies classified into quantile groups for experimental thermal tolerance metrics (i.e., CBRI Top20 ED50, CBRI Top20 ED5, CBRI Top20 DW, etc.). Due to heteroscedasticity and non-linear responses of coral colonies with varying thermal tolerance, a quantile regression model (package ‘*quantreg*’), was employed to robustly estimate the typical and extreme value of the bleaching response^60^, thereof revealing whether the examined thermal tolerance metrics better predict the bleaching outcomes for heat tolerant (i.e., lower CBRI quantiles) or heat susceptible colonies (i.e., higher CBRI quantiles). Hereby, quantile groups were determined a priori to mirror GLM model ranking, hence tau (τ) values were selected to approximate CBRI-based percentile groupings (τ = 0.2, 0.4, 0.6, and 0.8), and τ = 0.99 was used to specifically evaluate extreme bleaching outcomes within the upper tail of the distribution (i.e., full post-bleaching mortality). Accordingly, the quantile-specific relationship between experimental metrics (ED5, ED50, DW, ΔF_v/_F_m_, ΔBIS) and in situ bleaching metrics (i.e., CBRI in June and CBRI in October) was examined. Of note, any analysis including thermal stress metrics based on DRC modelling (i.e., ED50, ED5, DW), included only the colonies with a steady F_v_/F_m_ decline across treatments (see above). The remaining analyses (i.e., ΔF_v/_F_m_ and ΔBIS) included the full dataset (Table 1).

## Results

### Record-breaking heat stress and bleaching in Pulau Lang Tengah, Malaysia

Heat stress started to accumulate in early April 2024 and reached peak levels across all sites on June 18 (DHW = 9.6 °C-weeks) and June 21 (nDHW = 10.9 °C-weeks) 2024, respectively. Experienced heat stress levels thus exceeded the previous maximum level recorded in this region in 2010 by 4.4 °C-weeks (DHW_2010_ = 5.2) and 3.3 °C-weeks (nDHW_2010_ = 7.6 °C-weeks), respectively (Figure 1, Supplementary Figure 3-4). Importantly, heat stress levels were equally severe across sites; and although heat stress was severe across both depths, nDHW were on average 1.7 °C-weeks lower at 15 m depth than at 5 m (Supplementary Figure 5). Consequently, these heat stress levels resulted in severe bleaching as 92.2% of colonies of all species were bleached by June 2024 (mean CBRI June = 43.5), while 78.3% of colonies died partially, and 42.0% of colonies experienced full mortality by October 2024 (mean CBRI October = 53.6) (Figure 2). Supplementary Table 1 contains information on bleaching severity, mortality, and recovery.

**Figure 1.**
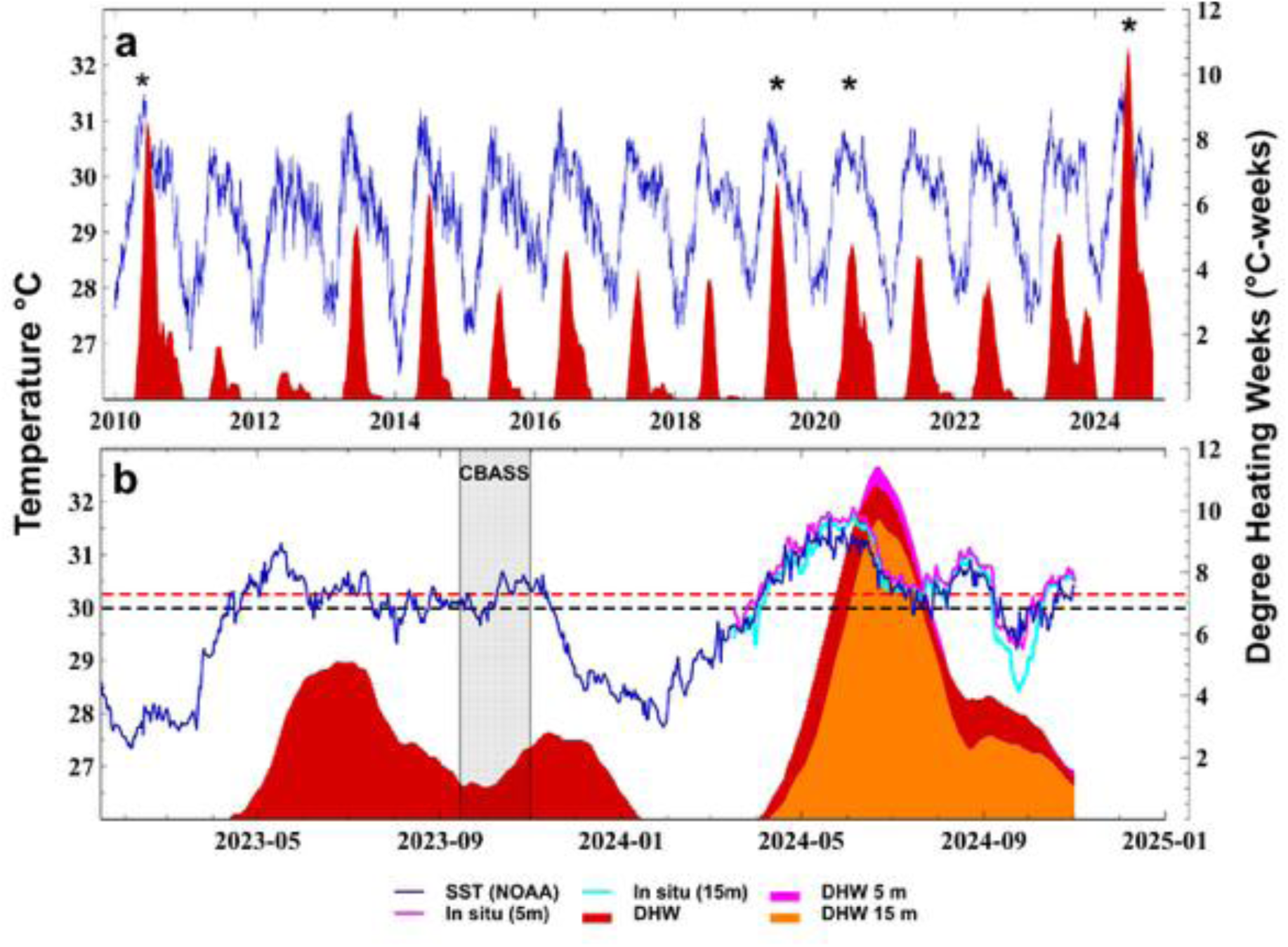
Record breaking heat stress in 2024, with (a) showing heat stress events since the previous record heat stress event in 2010 and (b) showing heat stress for the last 2 years during the study period, highlighting in situ heat stress across water depth. Blue lines show the satellite-measured sea surface temperatures (SST °C) between 1 January 2010 and 31 October 2024 around Pulau Lang Tengah (5.793844, 102.895492), recorded by the National Oceanic and Atmospheric Administration (NOAA), Coral Reef Watch (CRW) product, version 3.1. Heat stress is described as the accumulated daily mean temperature above the MMM, i.e., Degree Heating Weeks (DHW). A locally optimized DHW metric (i.e., sensu Szereday et al., 2024) is shown in dark red in both panels, based on temperature measured by NOAA satellites. Asterisks in panel (a) highlight heat stress events that resulted in coral bleaching (identified by available literature and author observations). The black-dotted line in panel (b) shows the historical maximum monthly mean (MMM) temperature (i.e., 29.94 °C) established by NOAA CRW. Panel (b) additionally shows available 24h daily in situ sea temperature measurements at 5 m (cyan trend line) and 15 m (blue trend line) water depth, respectively. The red-dotted line highlights the MMM (i.e., 30.24 °C) based on in situ temperature data from five sites around Pulau Lang Tengah recorded between 2020-2022 (see Szereday et al., 2024, 2025). DHW based on in situ temperature data is shown for both depths (DHW 5m and DHW 15m). The grey shaded area shows the 7-week period during which acute heat stress assays (i.e., CBASS) were conducted.

**Figure 2.**
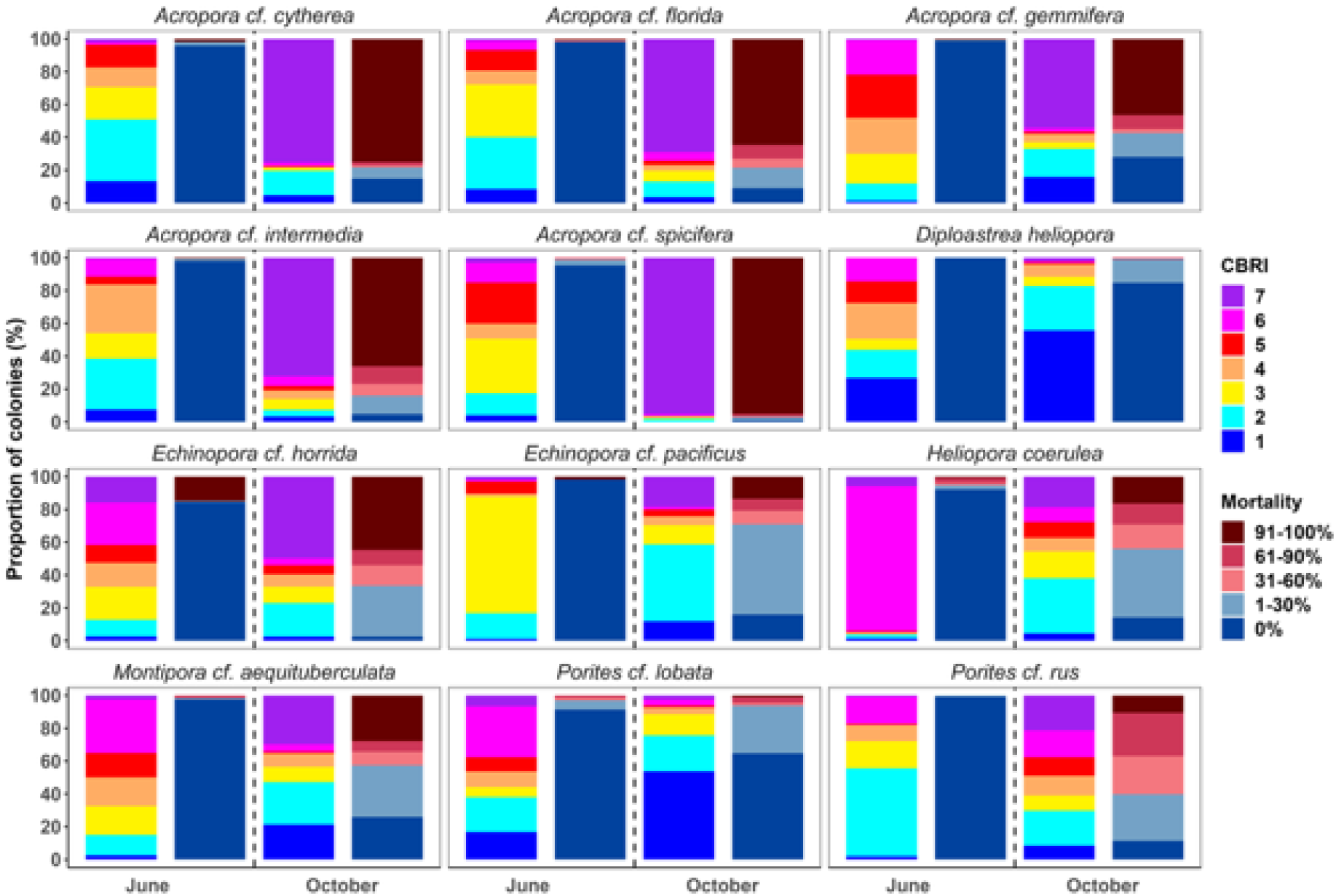
Coral bleaching severity and percentage of healthy, partially, and dead coral colonies in June 2024 during peak heat stress and four months later in October 2024. The severity of bleaching on a colony-level is expressed by the weighted Colony Bleaching Response Index (CBRI), whereby higher CBRI values (0-100) indicate more severe bleaching. The CBRI is presented together with the proportion of colonies with dead tissue for each time point, respectively, to distinguish the ecological interpretation of CBRI values across time points. High CBRI values in June primarily captured bleaching responses during peak heat stress, whereas CBRI values in October primarily reflect widespread post-bleaching mortality. The dashed line separates the two data points. CBRI categories: B1 – no bleaching, B2 ≤ 16.7, B3 =16.8-33.3, B4 = 33.4-50.0, B5 = 50.1-66.7, B6 = 66.8-83.3, B7 - CBRI > 83.3.

### Correspondence between standardized heat stress metrics and in situ bleaching

Prior to analysis, about half of all assayed colonies were removed from ED modelling due to violating the DRC model assumption (i.e., declining F_v_/F_m_ with increasing temperature) (Table 1). For the remaining colonies, we examined the correspondence of standardized experimental heat stress metrics (i.e., ED5, ED50, DW) with bleaching severity during peak heat stress (i.e., CBRI June) and with post-bleaching mortality four months later (i.e., CBRI October) for the same 999 and 992 colonies, respectively (sample sizes varied due to colony re-detection in situ).

We first tested for significant differences in thermal tolerance metrics between upper and lower quantile groups within each species to evaluate the discriminatory power of these metrics in distinguishing bleaching-tolerant from susceptible colonies, under the assumption that quantile-based grouping reflects true biological distributions. Using Dunn’s pairwise comparisons, significant differences between the Top20 vs. the Bottom20/Bottom40 and the Top40 vs. the Bottom20/Bottom40 were found for all species for each metric (BH-adjusted p ≤ 0.01, Supplementary Data 1). The distribution of thermal tolerance metrics is shown in the Supplement (Supplementary Figures 6-8), demonstrating a broad distribution of ED5, ED50, and DW values of the examined colonies.

To correlate experimental thermal tolerance metrics (ED5, ED50, and DW) with bleaching and post-bleaching mortality following record heat stress, the mean maximum likelihood probability of bleaching was estimated using general linear models for each colony, while accounting for environmental covariates. The predicted probabilities were extracted and visualized to identify species-specific patterns across metrices and time points (Figure 3 and 4). Of the 12 species examined, we found correspondence between higher ED5s with lower bleaching in June and post-bleaching mortality in October for *Diploastrea heliopora*, *Montipora* cf*. aequituberculata*, and *Echinopora* cf. *pacificus* (Figure 3 and 4). By comparison, lower bleaching and post-bleaching mortality corresponded with lower ED5 and ED50 values for *Acropora* cf. *gemmifera, Echinopora* cf. *horrida,* and *Acropora* cf. *intermedia,* as well as *M. aequituberculata* for the ED50 metric. Crucially, regardless of whether associations between in situ bleaching and CBASS metrics were positive or negative, bleaching probability patterns were consistent for *D. heliopora, M. aequituberculata* (for the ED5 metric), *A*. *gemmifera,* and *E*. *horrida*. Furthermore, inverted patterns were found when examining bleaching and post-bleaching mortality probabilities for the DW metric, as higher DWs corresponded to higher post-bleaching mortality for *D. heliopora*, *M. aequituberculata*, *E. horrida*, and *E. pacificus* (Figure 3). The same pattern was also observed when examining bleaching severity, except for *E. pacificus* (higher DW correlated with lower bleaching in the Top 20 quantile). Taken together, when positive correspondence between experimental and in situ bleaching metrics existed (i.e., higher experimental metrics = lower bleaching), these were found for ED5 and ED50 metrics for some species, with opposite trends for the DW metric.

**Figure 3.**
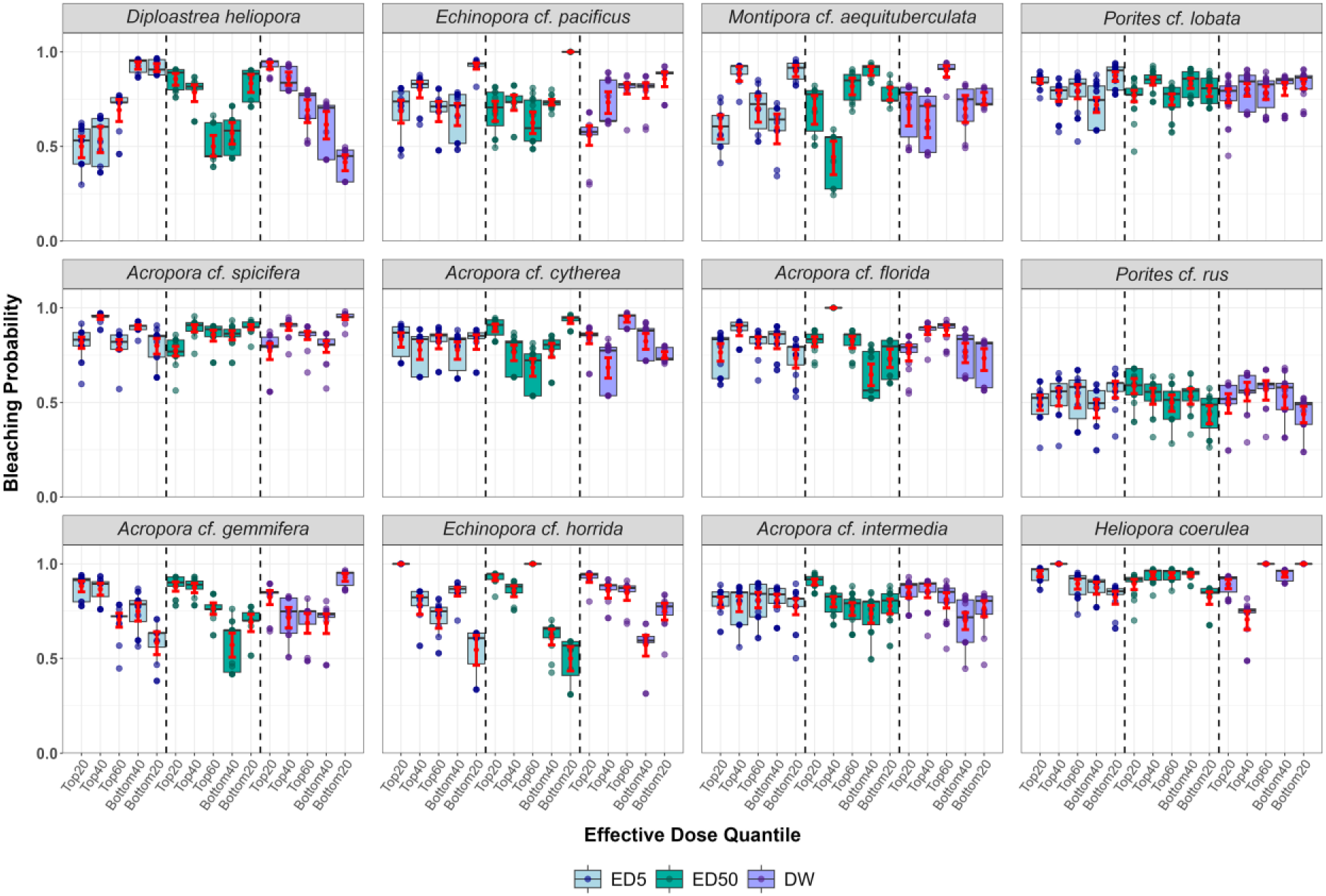
GLM-based estimation of in situ bleaching as a function of standardized experimental thermal tolerance metrics. ED5 signifies the onset temperature of thermal stress, ED50 the thermal tolerance threshold, and DW shows whether photosynthetic efficiency decline is gradual (higher DW) or more rapid (lower DW). Coral colonies are ranked based on ED-based quantiles (i.e., from Top20 to Bottom20). The model included ‘depth’ and ‘site’ as environmental covariates, and higher probabilities are representative of higher bleaching severity. Species are displayed in order of correspondence patterns: top row–correspondence in accordance with the initial hypothesis of lower bleaching and higher experimental thermal tolerance metrics; middle row–no discernible correspondence; bottom row–discrepancy between experimental and in situ metrics. This approach visualizes how the GLM estimated bleaching risk varies across colonies with low to high ED values. Red diamond squares and error bars show the mean and 95% bootstrapped confidence intervals. The median is shown by the bar plot. Dots represent individual data points. The dashed line separates ED5, ED50, and DW box plots for visual clarity.

**Figure 4.**
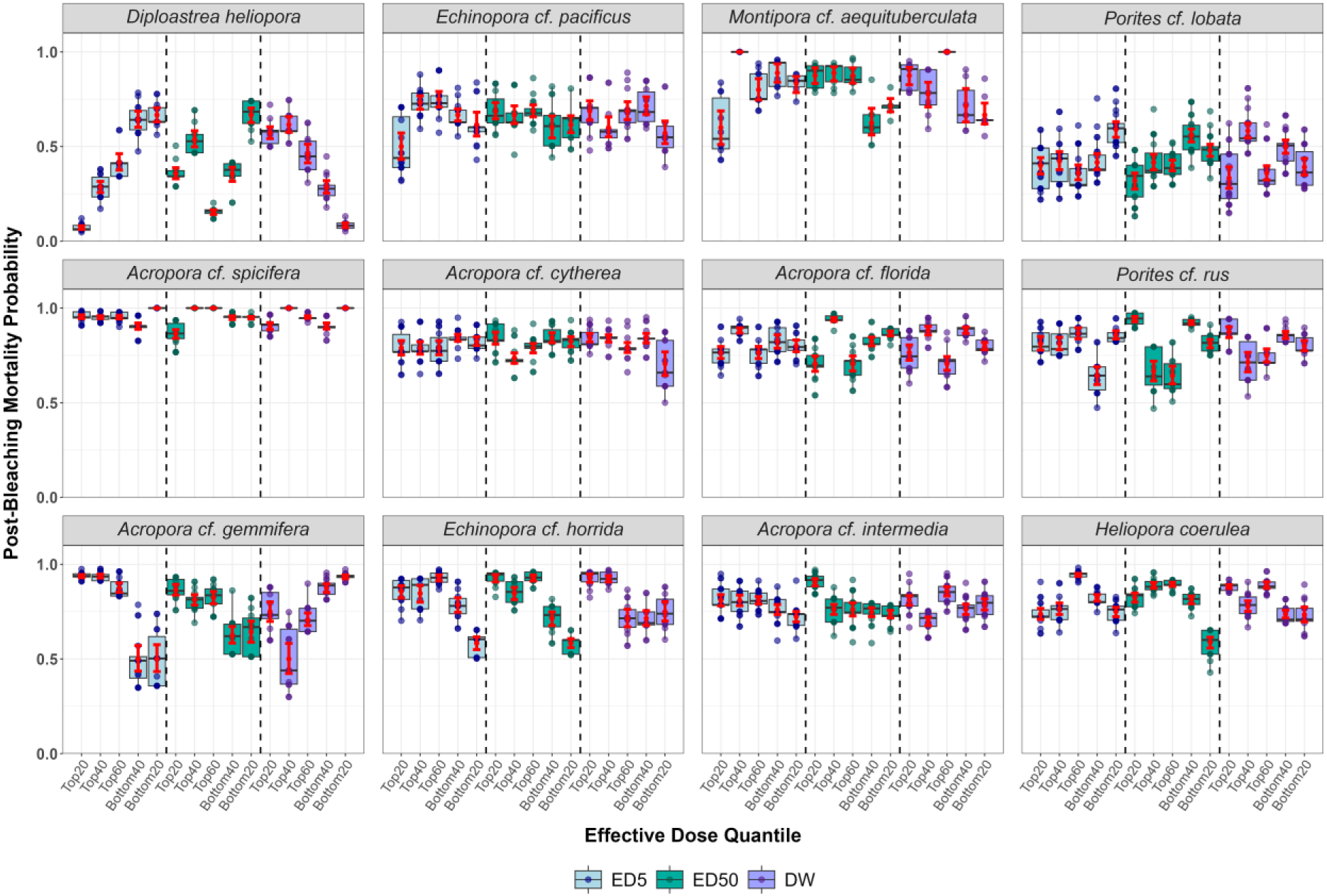
GLM-based estimation of in situ post-bleaching mortality as a function of standardized experimental thermal tolerance metrics. ED5 signifies the onset temperature of thermal stress, ED50 the thermal tolerance threshold, and DW shows whether photosynthetic efficiency decline is gradual (higher DW) or more rapid (lower DW). Coral colonies are ranked based on ED-based quantiles (i.e., from Top20 to Bottom20). The model included ‘depth’ and ‘site’ as environmental covariates, and higher probabilities are representative of higher post-bleaching mortality. Species are displayed in order of correspondence patterns: top row–correspondence in accordance with the initial hypothesis of lower bleaching mortality with higher experimental thermal tolerance metrics; middle row–no discernible correspondence; bottom row–discrepancy between experimental and in situ metrics. This approach visualizes how the GLM estimated mortality risk varies across colonies with low to high ED values. Red diamond squares and error bars show the mean and 95% bootstrapped confidence intervals. The median is shown by the bar plot. Dots represent individual data points. The dashed line separates ED5, ED50, and DW box plots for visual clarity.

To further explore these patterns, pairwise comparisons of CBRI values were conducted for quantile-ranked colonies, and mean group responses were examined using quantile regression. Pairwise comparisons using Dunn’s test revealed limited significant correspondence between in situ bleaching and experimental thermal tolerance across species and time points. Significant mean post-bleaching mortality differences (i.e., October CBRI) were only found for *D. heliopora* for the ED5 metric (Figure 5). Specifically, the mean CBRI of Top20 colonies was significantly lower than that of colonies classified within the Bottom20 and Bottom40 quantiles (pairwise Dunn’s test, p=0.016 vs. Bottom20 and p=0.024 vs. Bottom40). As found with GLMs, lower DWs were significantly correlated with in situ bleaching for *D. heliopora.* Despite this, Top20 ranked colonies were significantly more bleached and dead than Bottom20 colonies in June and October (p=0.013 in June and p=0.043 in October), respectively. No additional significant associations were detected between higher ED50, ED5, or DW values and reduced in situ bleaching (Supplementary Figures 9-10, Supplementary Data 2).

**Figure 5.**
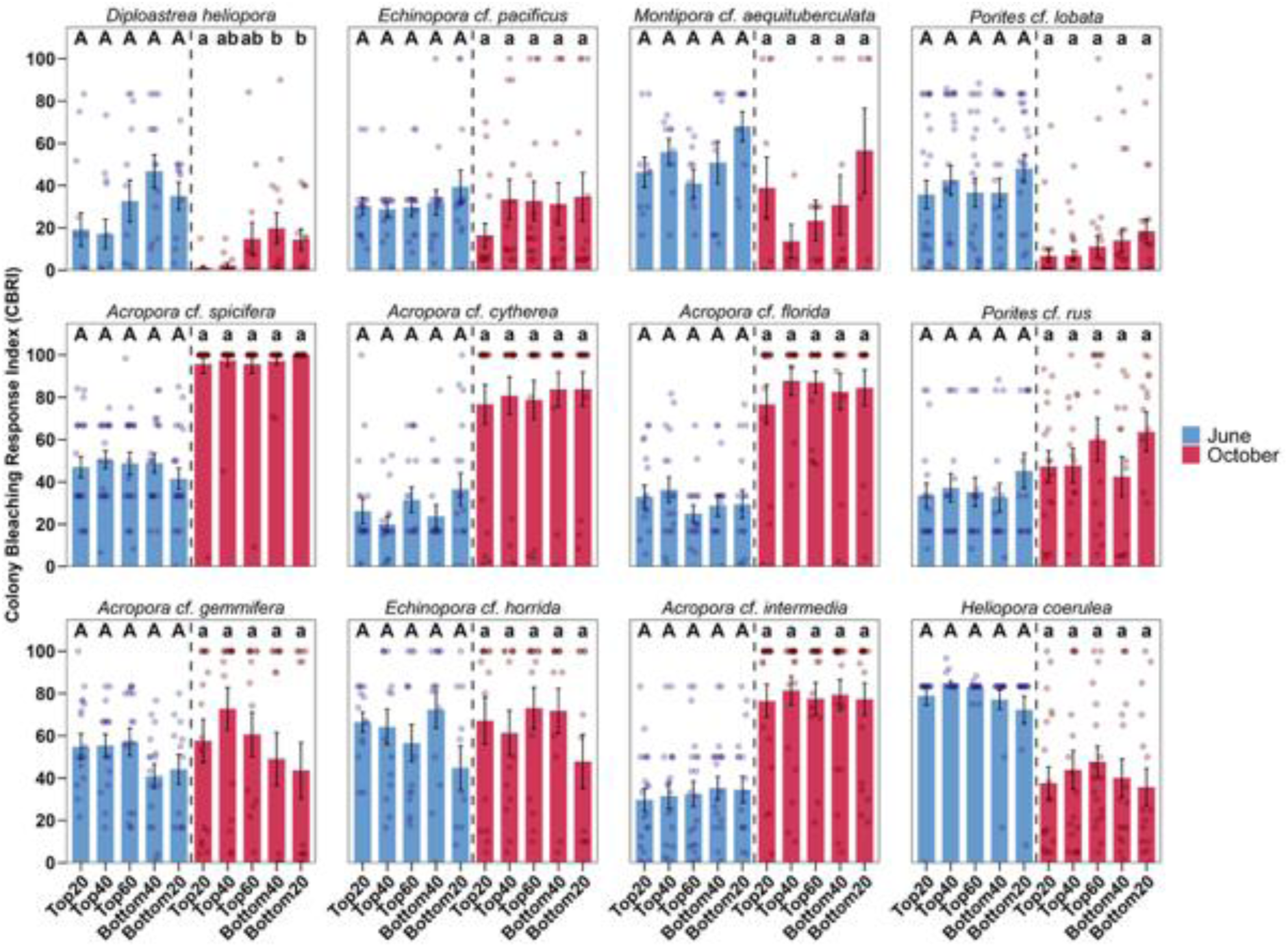
Comparison of bleaching severity and post-bleaching mortality between coral colonies in the upper and lower ED5 quantiles. Coral bleaching outcomes (i.e., Colony Bleaching Response Index) were measured in June 2024 during peak heat stress and four months later in October 2024. These measurements capture the immediate heat stress response and subsequent mortality, respectively. ED5 is the thermal breakpoint temperature, i.e., the temperature signifying the onset of thermal stress, i.e., the onset of a decline in photosynthetic efficiency. Error bars signify the standard error (±), and dots represent individual CBRI data points of each coral colony. Annotated letters alternate in each panel between upper (i.e., June) and lower case (i.e., October) letters for clarity. Letters represent results of Dunn’s Test conducted at significance levels of p ≤ 0.05.

In a final step, because GLM-derived probabilities and pairwise comparisons did not fully align, we applied quantile regression to estimate bleaching and post-bleaching mortality across conditional quantiles of the response as a function of experimental thermal tolerance metrics. Quantile regression was applied following GLMs and pairwise comparisons to evaluate how experimental thermal tolerance relates to in situ bleaching severity and post-bleaching mortality across the response distribution, testing whether these metrics are more informative for predicting extreme versus mild outcomes. We regressed the bleaching severity of 999 colonies (i.e., CBRI June) and post-bleaching mortality of 992 of the same colonies (i.e., CBRI October) against the ED5, ED50, and DW metrics in separate model iterations. Overall, we detected both positive (higher experimental thermal tolerance associated with increased bleaching) and negative (higher tolerance associated with reduced bleaching) relationships across species and metrics, although these patterns were inconsistent across species and quantiles, pointing to unresolved confounding factors (e.g., species traits, algal assemblages, or environmental heterogeneity) (Table 2).

**Table 2.**
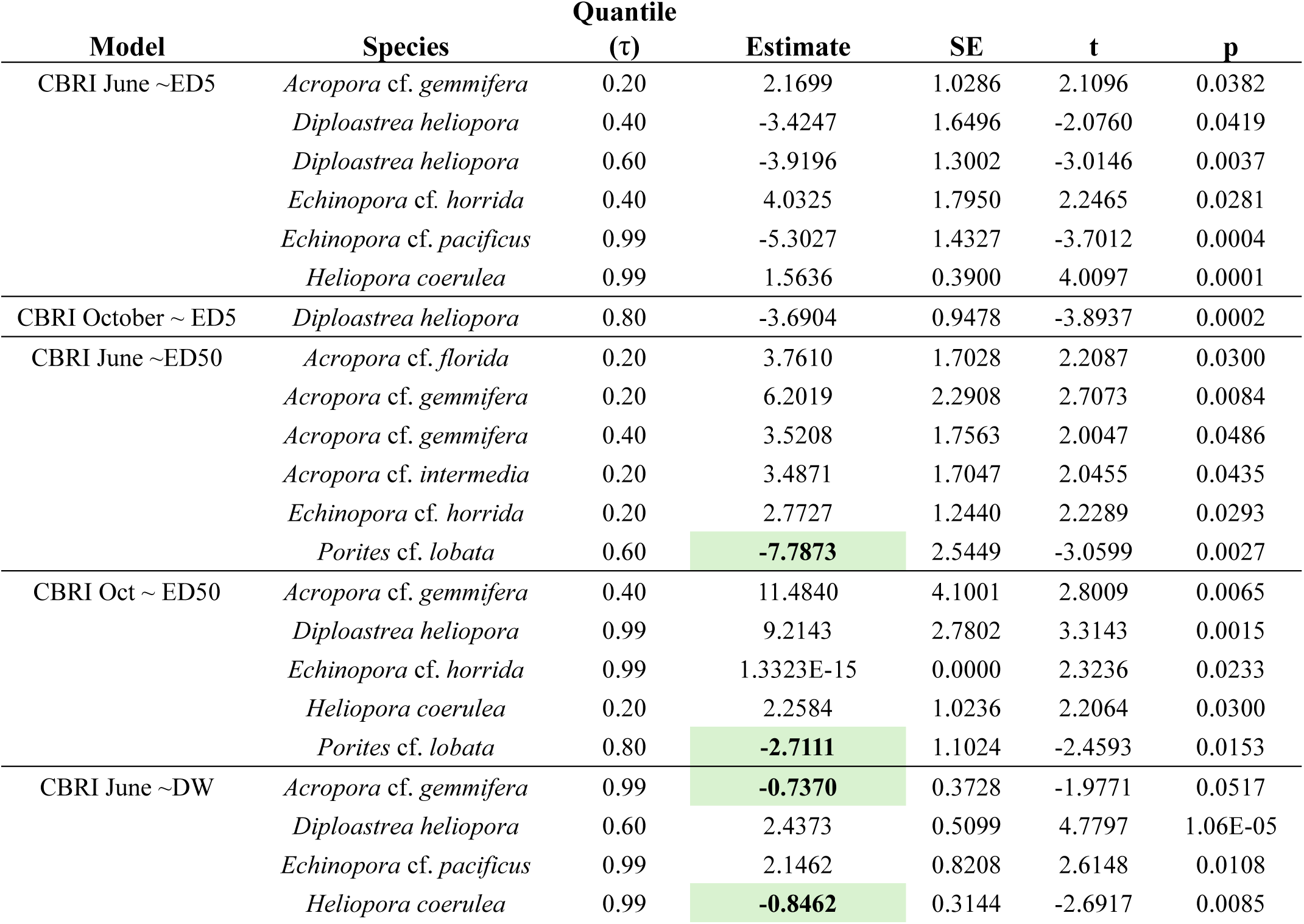
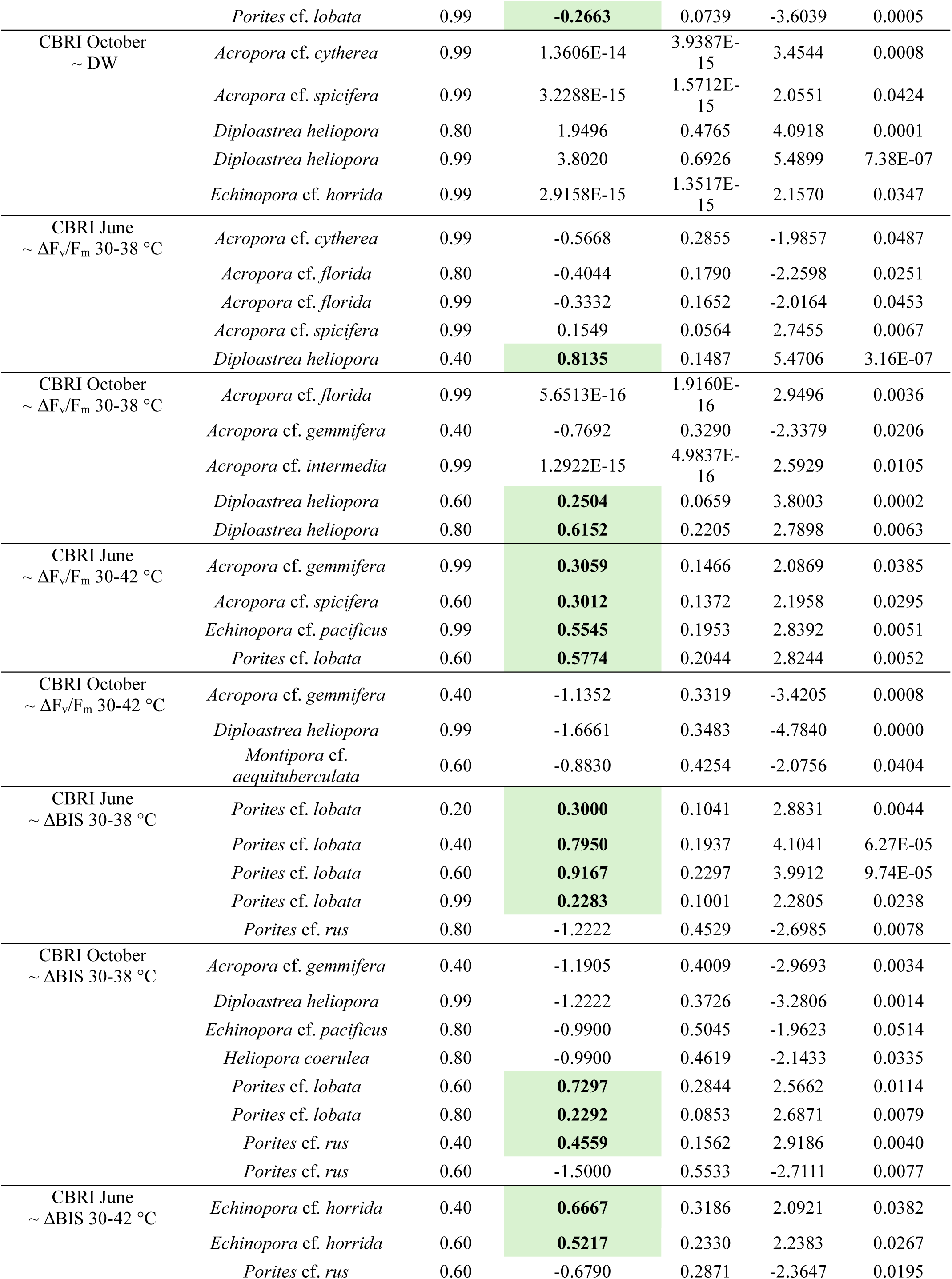
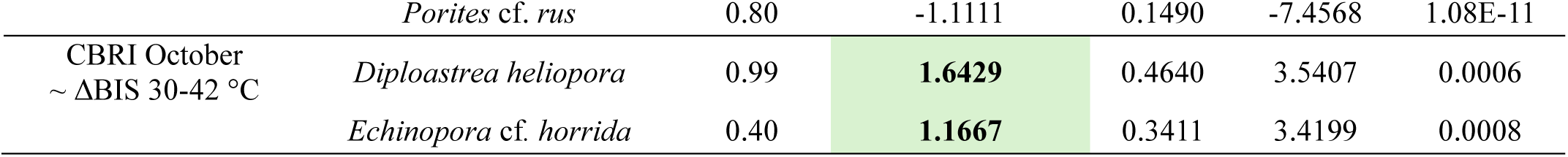
Summary of significant quantile regression results with non-zero estimates for bleaching severity and post-bleaching mortality. Coral bleaching severity was measured in June 2024, and post-bleaching mortality was measured in October 2024. Higher bleaching severity and mortality are reflected by higher Colony Bleaching Response Indexes (CBRI) measured at the respective time point. Colonies were ranked into top and bottom quantiles (i.e., *τ* = 0.20, 0.40, 0.60, 0.80, 0.99) based on their CBRI values. Quantile regression was conducted for each time point and for each experimental thermal tolerance metric (i.e., ED5, ED50, DW) in individual iterations. CBRI values were regressed against standardized thermal tolerance metrics: ED50–thermal tolerance threshold at which 50% initial photosynthetic efficiency is decreased by 50%; ED5–breakpoint temperature at which the decline in photosynthetic efficiency is initiated; DW calculated as ED95-ED5, with ED95 being the thermal limit at which photosynthetic efficiency is decreased by 95%, whereby higher values are suggestive of increased thermal resilience. ΔFv/Fm is the change of photosynthetic efficiency between the highest (i.e., 38 °C and 42 °C) and baseline (i.e., 30 °C) temperature treatments (i.e., higher values reflective of a more severe decline); ΔBIS was calculated in a similar manner based on visual Bleaching Index Scores (BIS). SE – standard error. Bold highlights cases for which CBASS-based metrics significantly corresponded with lower in situ bleaching outcomes.

Crucially, quantile regression recapitulated patterns identified by GLMs. For example, for *D. heliopora,* higher ED5 values were significantly associated with lower post-bleaching mortality of the Bottom40 quantile colonies (i.e., τ = 0.80, p = 0.0002), while higher ED5 values were significantly associated with lower bleaching severity in the Top40 (p=0.0419) and Top60 (p=0.0037) quantile colonies, corroborating GLM and pairwise Dunn’s Test results (Figures 3 and 5). Furthermore, counter to the initial hypothesis, GLMs identified positive associations between higher ED5 and higher bleaching severity (Figure 3), which were corroborated by quantile regression in Top20 *A. gemmifera* (p=0.0382), Top40 *E. horrida* (p=0.0281), and Bottom20 *H. coerulea* (p=0.0001) (Table 2). This positive associations was particularly pronounced for the ED50 metric, as higher bleaching severity with higher ED50 values were detected for Top20 (p=0.0084) and Top 40 *A. gemmifera* (p=0.0486), Top20 *A. intermedia* (p=0.0435), Top20 *A. florida* (p=0.03), and Top20 *E. horrida* (p=0.0293), while in contrast lower bleaching associated with higher ED50 in Top60 *P. lobata* colonies (p=0.0027), and lower post-bleaching mortality in Bottom40 *P. lobata* colonies (p=0.0153). Moreover, consistent with GLM models and pairwise Dunn’s test, higher post-bleaching mortality with increasing DW was found for *D. heliopora* colonies in the lower Bottom40 and Bottom20 quantiles (p<0.001), suggesting that among the most susceptible colonies, those with higher DW experienced higher post-bleaching mortality. Collectively, these findings reveal pronounced interspecific differences in the relationship between experimental thermal tolerance metrics and bleaching responses.

To leverage the full dataset beyond dose–response curve modelling constraints (Table 1), we quantified relative changes in F_v_/F_m_ and BIS between the baseline and the two highest temperature treatments (i.e., ΔF_v_/F_m_ 38°C, ΔBIS 38°C, ΔF_v_/F_m_ 42°C, ΔBIS 42°C). These metrics were then modelled as a function of CBRI for all colonies subjected to heat stress assays and surveyed for bleaching severity (i.e., 1,863 colonies in June) and post-bleaching mortality (i.e., 1,853 colonies in October). Herein, similar relationships as for DRC-based metrics were found. Consistent with the initial hypothesis that greater declines in ΔF_v_/F_m_ with increasing temperature correspond to higher bleaching susceptibility, larger ΔF_v_/F_m_ between 30°C and 38°C were associated with increased post-bleaching mortality in *D. heliopora* within the Top60 (p = 0.0002) and Bottom40 (p = 0.0063) quantiles. However, estimated effect sizes were modest (0.2504 and 0.6152, respectively). In contrast, inverse relationships were observed, with lower post-bleaching mortality despite higher ΔF_v_/F_m_. This pattern was evident for Top40 *A. gemmifera* (ΔF_v_/F_m_ 38°C: p = 0.0206; ΔF_v_/F_m_ 42°C: p = 0.0008), consistent with GLM and DRC-based predictions for this species. Similar inverse associations were also detected for *M. aequituberculata* (Top60, p = 0.0404) and *D. heliopora* (Bottom20, p < 0.0001) for ΔF_v_/F_m_ at 42°C.

Visual assessment of pigmentation changes (i.e., ΔBIS) of experimental fragments with increasing temperature provided limited predictive insight (Table 2). Higher values of the experimental metric ΔBIS 30-38 °C were significantly associated with higher in situ bleaching severity in June for *P. lobata* colonies in the Top20 (p=0.0044), Top40 (p<0.0001), Top60 (p<0.0001), and Bottom20 (p=0.0238) quantiles (Table 2). These associations were partially recaptured when assessing post-bleaching mortality for Top60 and Bottom40 *P. lobata* in October 2024 (p=0.0114 and 0.0079, respectively) (Table 2). However, when regressing against the ΔBIS 30-42 °C metric for *P. lobata*, no significant relationships were found, suggesting that relative BIS change at the highest temperature treatment is not a differentiating factor for this species. Significant higher post-bleaching mortality associated with higher ΔBIS 30-42 °C for *E. horrida* (Top40, p=0.0008) and *D. heliopora* (Bottom20, p=0.0006). No additional associations supported the initial hypothesis, indicating that these experimental metrics offer limited ability to reliably predict in situ bleaching tolerance across different coral species. Full statistics tables are provided in the Supplementary Data 3.

## Discussion

Coral reefs worldwide continue to decline, necessitating urgent advances in conservation and restoration strategies. In this context, there is growing interest in improving the rapid identification of bleaching-resistant and resilient coral colonies. However, it remains poorly understood how well experimentally derived thermal tolerance metrics predict in situ bleaching severity and mortality, particularly under severe heat stress. To address this gap, we evaluated the correspondence between commonly used experimental thermal tolerance metrics and field-based bleaching severity and post-bleaching mortality across 999 and 992 coral colonies, respectively. During the study period, accumulated heat stress reached record levels, resulting in partial mortality of 78.3% across surveyed colonies, and complete mortality of 42.0% of surveyed colonies, largely independent of depth and reef site^19^. Accordingly, the 2024 bleaching event represents the most severe event documented in northeastern Peninsular Malaysia to date^19,46,49,61,62^.

Following record heat stress exposure, we found correspondence between at least one of the examined experimental thermal tolerance metrics (ED5, ED50, DW) determined prior to heat-loading with bleaching and post-bleaching mortality in the wild for three species (i.e., *D. heliopora*, *M. aequituberculata*, and *E. pacificus*), no discernible correspondence for six species (*A. spicifera*, *A. cyhtherea*, *A. florida*, *H. coerulea*, *P. rus*, and *P. lobata*), and inverted correspondence for three species, i.e., higher bleaching and mortality despite higher experimental thermal tolerance (*A*. *gemmifera, E. horrida,* and *A. intermedia*). Thus, correspondence between experimental and in situ responses was strongly species- and metric-dependent, with the main patterns identified by the GLMs being largely recaptured by the quantile regression analyses. For instance, higher thermal breaking point values significantly corresponded with lower mortality for *D. heliopora* under record heat stress, and comparable patterns were observed in other species, corroborated by quantile regression analyses. This suggests that thermal breaking points more closely reflect in situ bleaching susceptibility for some but not all species, echoing conclusions that even the most resilient corals may have limited capacity to tolerate heat stress beyond their current individual breaking points^27^. Conversely, other species exhibited weak, absent, or even inverted associations. Similarly, different metrics (e.g., ED5, ED50, DW, ΔF_v_/F_m_) did not perform equivalently, and in some cases yielded opposing relationships with bleaching outcomes, the origin of which remains currently unresolved. Using quantile regression to model the relationship between experimental visual bleaching (ΔBIS 30–38 °C and ΔBIS 30–42 °C) and in situ bleaching, we found limited significant associations with lower in situ bleaching and post-bleaching mortality. Where present, significant relationships were found in slow-growing species and were not consistent across quantiles. Thus, discrepancies across metrics existed across species regardless of whether assessed visually (BIS) or by means of algal fluorometry (F_v_/F_m_). Taken together, while significant associations between experimental metrics and field outcomes were detected for some species and metrics, these relationships were inconsistent in direction and strength across taxa and quantiles.

Two things should be considered when interpreting these results: the level of resolution sought and the environmental context. Acute heat stress assays such as CBASS were developed to provide rapid, standardized estimates of coral thermal tolerance under field conditions^33,35,36^, and prior studies have demonstrated their ability to resolve ecologically meaningful differences across regions, reef sites, and populations^33,45^. The present study addressed a more stringent and underexplored question: whether experimentally derived thermal tolerance metrics can resolve differences among colonies within populations and predict subsequent bleaching severity and post-bleaching mortality in situ. Specifically, we tested the ability to resolve variation among colonies within populations using a fine-grained quantile framework (20% quantiles), which represents a substantially more demanding application. Our results indicate that this was only partially successful. However, rather than a binary validation or rejection of acute heat stress assays, our findings point to a more nuanced conclusion: the predictive value of experimentally derived thermal tolerance is conditional on species, metric choice, and environmental context. Thus, it must be acknowledged that the discriminatory power of such assays decreases as the scale of inference shifts from population-level contrasts to within-population ranking. In other words, differences between populations are easier to detect because many colony replicates provide a robust mean for comparison, whereas within-population, colony-level assessments rely on single measurements per colony. These findings caution against interpreting experimental thermal tolerance metrics as direct proxies for bleaching susceptibility without considering species-specific and ecological contexts.

Regarding the environmental context, the 2024 heat stress event in northeastern Peninsular Malaysia was extreme and record-breaking by any means, with widespread bleaching and substantial mortality across species. Under such conditions, bleaching responses may converge, reducing the observable separation among colonies that differ in experimentally derived thermal tolerance. It should also be noted that roughly half of the colonies did not exhibit decreasing F_v_/F_m_’s with increasing temperature, suggesting heterogeneous photo-physiological responses, which remain unexplained at this point. Previous studies suggest that acute heat stress assays may accurately differentiate thermal tolerance primarily under moderate in situ heat stress^61^. This notion is supported by field data from the Great Barrier Reef, which found a predictive relationship between experimental visual bleaching assessments and higher bleaching survival in situ^63^. Conversely, under extreme heat stress in 2023, almost all genotypes of *Acropora* species across the Florida Reef Tract were equally affected, regardless of ascribed thermal tolerance capacity^64^. Similarly, field studies from all ocean basins have shown high within-species mortality following record heat stress^6,17,19,65,66^.

Considering the above, our results–in line with previous studies–indicate that under extreme heat stress, bleaching and mortality responses may converge, reducing the extent to which differences in experimentally derived thermal tolerance and underlying genetic variation are expressed. This reduces the discriminatory power of acute heat stress assays under such conditions, with implications for their application in conservation and restoration. It is, however, equally important to note that a small subset of colonies (approximately 5–10%) across 11 of the 12 species (excepting *A. spicifera*) did not bleach nor suffer mortality or only bleached to a modest extent and recovered therein^37^. Therefore, highly resistant individuals were found in situ, albeit these colonies were not consistently predicted by experimental metrics. These colonies now provide time-sensitive research opportunities to elucidate the underlying mechanisms of their extraordinary resilience.

From an applied perspective, our results have important implications for the use of acute heat stress assays in conservation and restoration. CBASS remains a useful tool for rapidly assessing thermal response and identifying broad patterns of variation across populations and environments. However, its use for fine-scale ranking of colonies within populations should be approached with caution, particularly when decisions depend on predicting long-term performance under natural conditions. A more robust approach is to integrate acute assay-derived metrics with complementary information, including longitudinal field observations, species-specific traits, and environmental context. Such integration is likely necessary to capture the multidimensional nature of coral resilience. Indeed, acute heat stress assays in isolation may not capture the complexity of in situ thermal tolerance and the bleaching response, which is influenced by the coral microbiome^65^, coral nutritional status^67^, and prevailing abiotic factors besides temperature (e.g., dissolved oxygen, salinity, water flow)^54,68^. Furthermore, experimental thermal tolerance metrics represent a momentary performance assessment, and repeated measurements over time may be required to obtain more consistent estimates of thermal tolerance that match real-world observations^67^. Our results further emphasize that selecting corals based solely on current bleaching tolerance may be insufficient, highlighting the need for approaches that actively enhance thermal tolerance^69^. Recognizing these constraints is essential to guide robust decision-making in the design, implementation, and interpretation of conservation and restoration strategies in coral reef research and management.

## Conclusion

Acute heat stress assays have emerged as useful tools for rapidly quantifying coral thermal tolerance under field conditions, but their ecological interpretation requires greater nuance than is often assumed. By comparing experimentally derived thermal tolerance metrics with subsequent bleaching severity and post-bleaching mortality across 12 Indo-Pacific coral species during the most severe heat stress event recorded in northeastern Peninsular Malaysia, this study shows that single acute assay outputs only partially capture realized bleaching trajectories in the wild. Correspondence between experimental and in situ responses was strongly species- and metric-dependent, with some species showing meaningful agreement, whereas others showed weak, absent, or even inverted relationships. Importantly, a small subset of colonies across most species remained resistant or recovered despite extreme heat stress, indicating that exceptional resilience persists in situ even when it is not consistently predicted by experimental screening. These colonies represent critical opportunities for further research and downstream interventions to expand the natural adaptive capacity of coral holobionts. Overall, our results support the continued refinement and use of acute heat stress assays as informative comparative and screening tools, but caution against treating them as universal stand-alone predictors of colony-level in situ performance. Effective application in conservation and restoration will require integration with species biology, environmental context, and fate-tracking corals in the field, i.e., longitudinal ecological validation.

## Supporting information

Electronic Supplementary Material (ESM)

## Author contribution

SS conceived the study. SS, CRV – conceptualized the study and manuscript; SS, KLC, JAH, CRV designed the ex-situ experiments. SS – designed coral bleaching surveys; SS, KLC, JAH, NZ conducted the heat stress experiments; SS and KLC collected the bleaching data: SS analysed the data and wrote the original draft; all authors edited and revised the original draft.

## Acknowledgement

Research was conducted under permit number Prk.ML.630-7Jld.12 (9) and Prk.ML.630-7Jld.14 (1) issued by the Department of Fisheries (DoF) Malaysia (Jabatan Perikanan Malaysia), and permit number EPU 40/200/19/3717 (11), issued by the Ministry of Economy (Kementerian Ekonomi) Malaysia. Our gratitude is extended to Chong Tze Seng and Sau Ying Chew from Summer Bay Resort Lang Tengah Island, and the entire dive centre crew for their continued support of field research activities. Thank you to our research assistants, Febrianne Sukiato, Kiu Yee Tong, and Dharshinie Mano for assisting with field work. We acknowledge lab technicians Kris and Miriam for determining visual bleaching scores.

