## Supplementary material for "Ecological bleaching trajectories under severe heat stress are only partially captured by acute heat stress assays": Electronic Supplementary Material (ESM)

Electronic Supplementary Material accompanying the article:

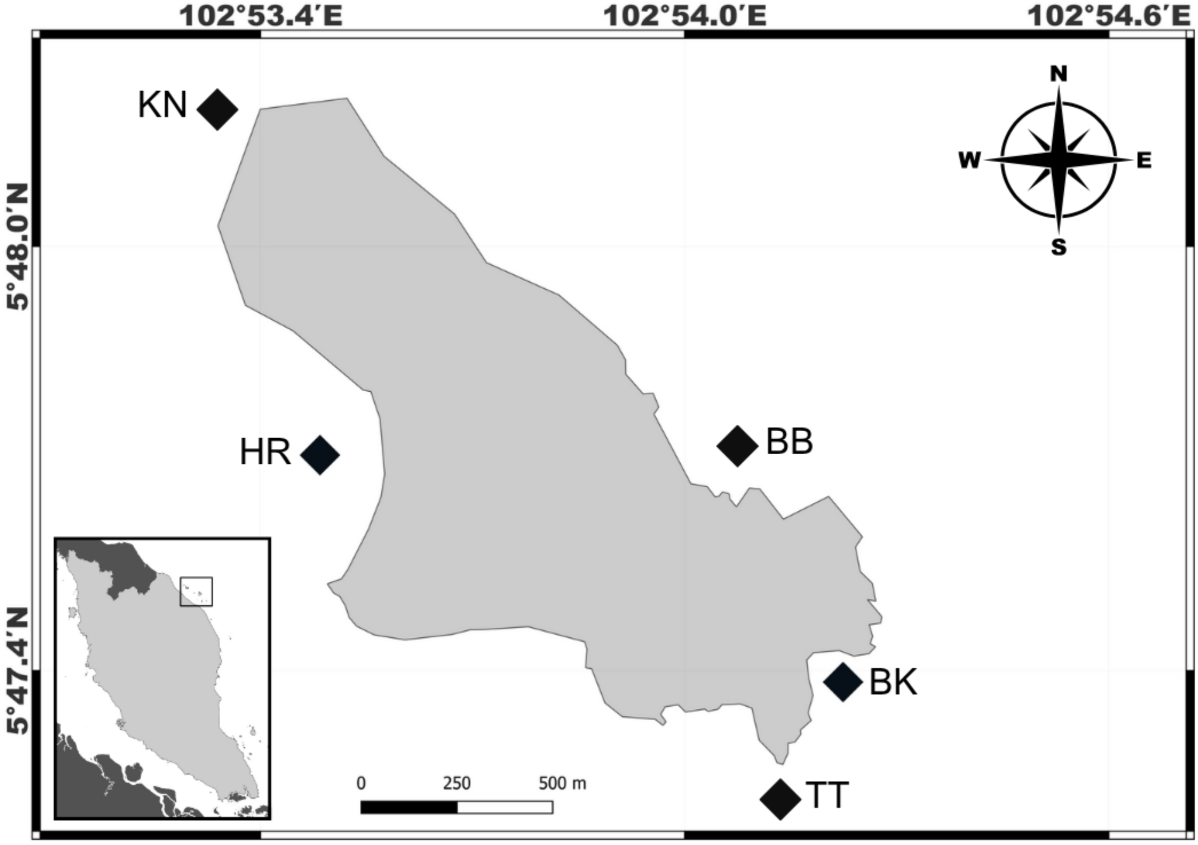


**Supplementary Figure 1**. **Geographic location of study sites around Pulau Lang Tengah (5.793844, 102.895492) along the northeastern coast of Peninsular Malaysia (zoomed out panel).** Site abbreviations BB – Batu Bulan, BK – Batu Kucing, TT – Tanjung Telunjuk, HR – House Reef, KN – Karang Nibong.

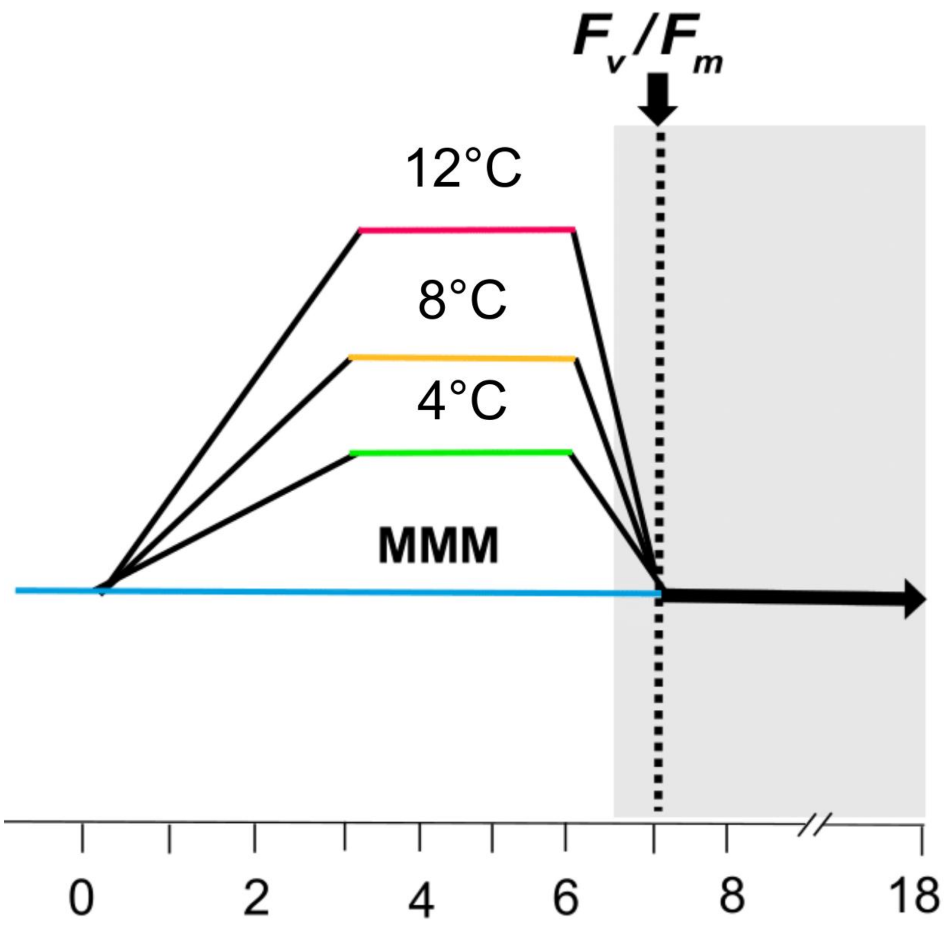


**Supplementary Figure 2.** Acute heat stress assay profiles of Coral Bleaching Automated Stress System (CBASS) with respective 3h maximum temperature treatments at 30 °C (baseline = maximum monthly mean MMM), 34 °C, 38 °C, and 42 °C. Maximum photosynthetic efficiency of PSII ($F_v/F_m$) of the coral fragments were measured after a 1-h dark acclimation as indicated by the grey shading.

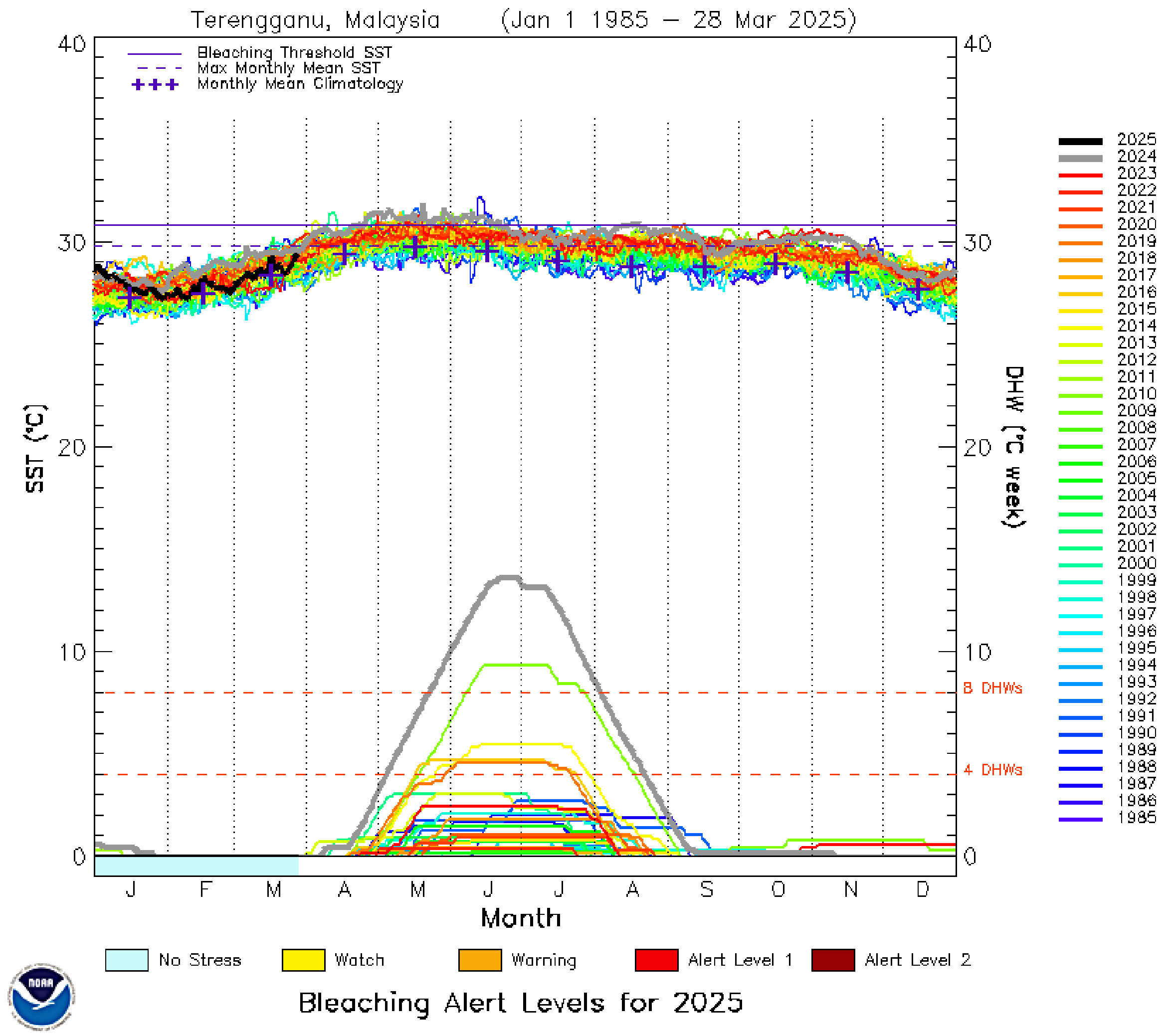


**Supplementary Figure 3.** Sea surface temperatures (SST; top) and cumulative heat stress degree heating weeks (DHW; bottom) data for Terengganu state, northeast Peninsular Malaysia, recorded by NOAA Coral Reef Watch (CRW) since 1985. Grey lines show the values through the 2024 heat stress events; coloured lines show each year's data from 1985-2023. Figure from NOAA Coral Reef Watch. Figure obtained from the CRW website at: https://coralreefwatch.noaa.gov/product/vs/gauges/terengganu.php

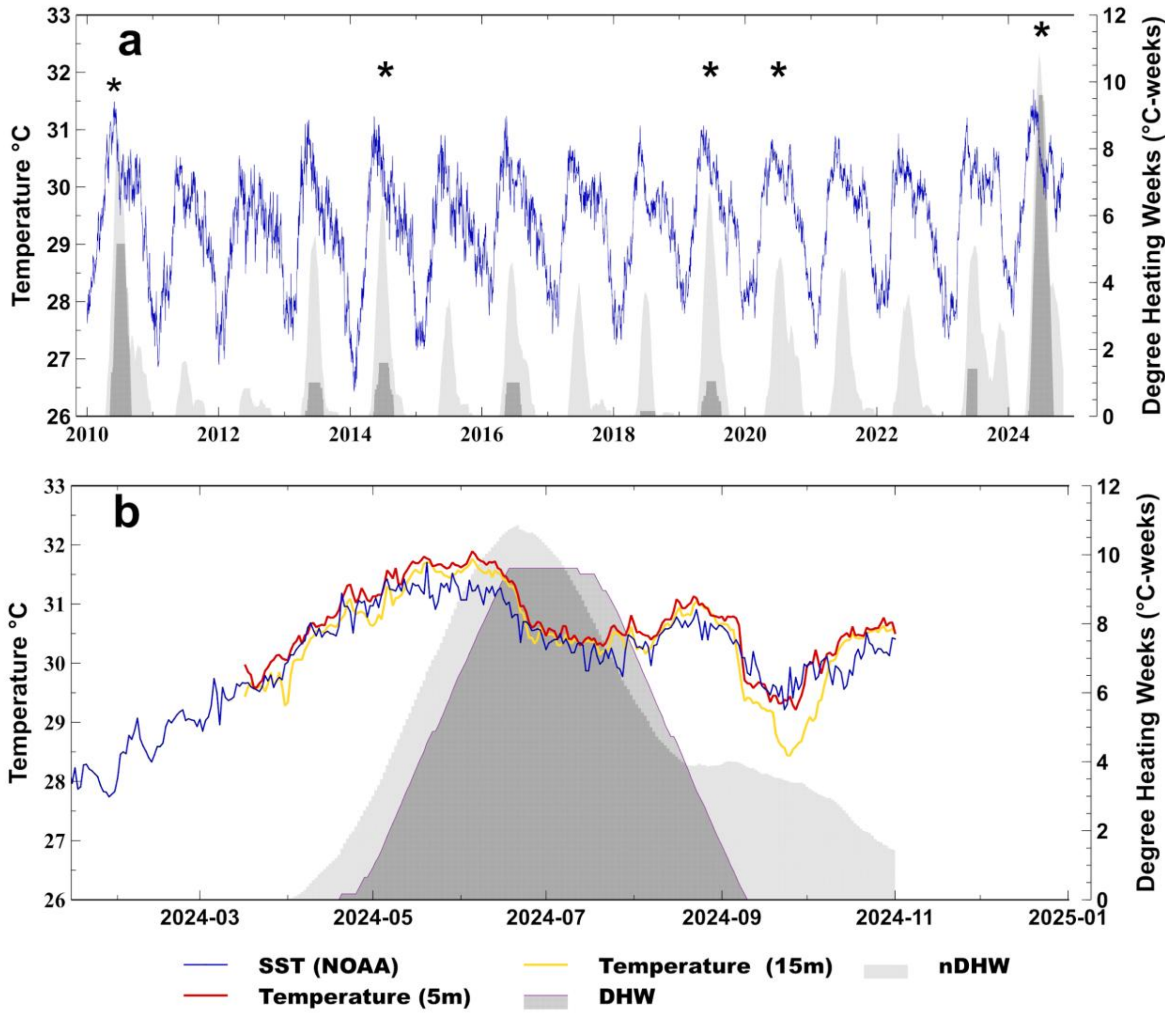


**Supplementary Figure 4**. **Sea surface temperature (SST) and degree heating weeks (DHW).** In both panels**,** the blue line shows the satellite-based nightly SST (°C) for Pulau Lang Tengah (5.793844, 102.895492), recorded by the National Oceanic and Atmospheric Association (NOAA), Coral Reef Watch (CRW) product, version 3.1. Panel (a) shows heat stress events since 2010 with observed coral bleaching events (identified by available literature and author observations) highlighted by an asterisk (*). SSTs are plotted against the standard DHW metric (dark-grey area), the adjusted nDHW metric (light-grey area), and the maximum monthly mean (MMM) temperature based on CRW satellite data (black dotted line, 29.94°C). Panel (b) shows average nightly satellite-based (blue line), in situ temperature measurements at 5 m (red line), and situ temperature measurements at 15 m (yellow line) during the 2024 record heat stress event.

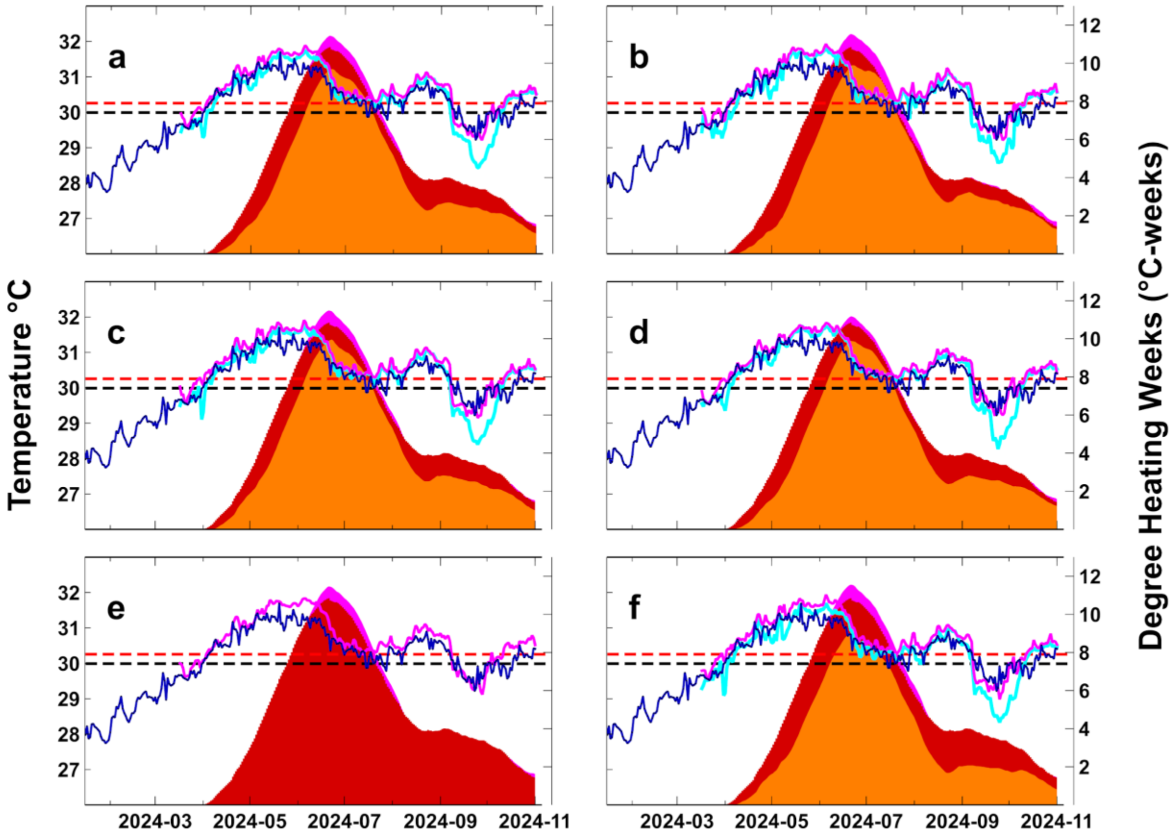


**Supplementary Figure 5. Sea surface temperature (SST) and degree heating weeks (DHW).** The blue line shows the satellite-based nightly SST (°C) between 1 January 2010, and 31 October 2024, around Pulau Lang Tengah (5.793844, 102.895492), recorded by the National Oceanic and Atmospheric Association (NOAA), Coral Reef Watch (CRW) product, version 3.1. The black-dotted line shows the maximum monthly mean (MMM) temperature based on NOAA CRW satellite data (i.e., 29.94 °C). The red-dotted line highlights the MMM (i.e., 30.24 °C) based on in situ temperature data from five sites around Pulau Lang Tengah recorded between 2020-2022 (see Szereday et al., 2024 for methodology). Heat stress based on satellite SST data (i.e., DHW NOAA) is shown in dark red. Panel (a) highlights average SST and in situ temperature at island scale for Pulau Lang Tengah, together with DHW based on satellite and in situ measurements (magenta = DHW at 5m, orange = DHW at 15 m). Panels (b-f) show site-specific temperature and heat stress data, where mean daily 24h in situ sea temperature are shown for measurements at 5 m (cyan line) and 15 m (blue line) water depth, respectively. Panel (b)- Batu Bulan; (c) – Batu Kucing; Panel (d) – House Reef; (e) – Karang Nibong (deep data missing due to logger malfunction); (f) – Tanjung Telunjuk.

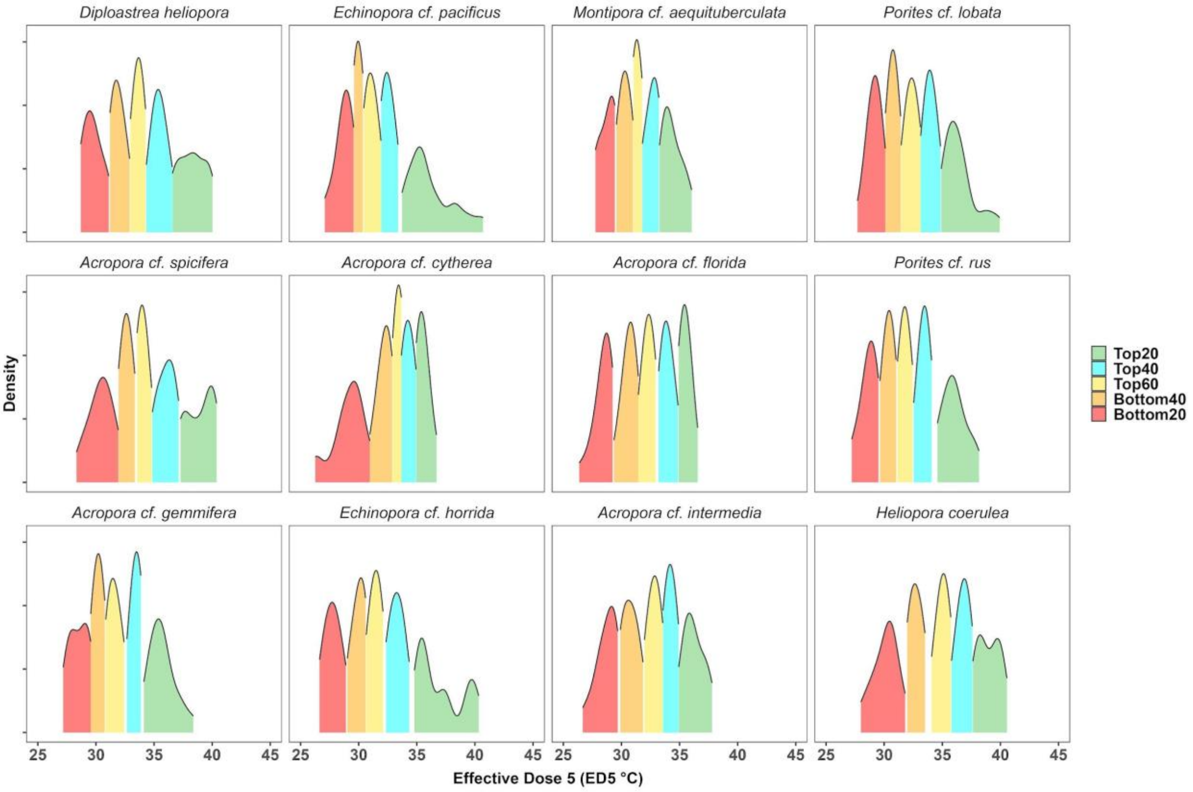


**Supplementary Figure 6. Species-specific density distributions of thermal breakpoint temperature (Effective Dose 5, ED 5 in °C) across coral colonies ranked into Top to Bottom quantile groups.** Thermal tolerance declines from top-ranked (green and cyan) to bottom-ranked (orange and red) corals, showing a clear stepwise pattern across all species.

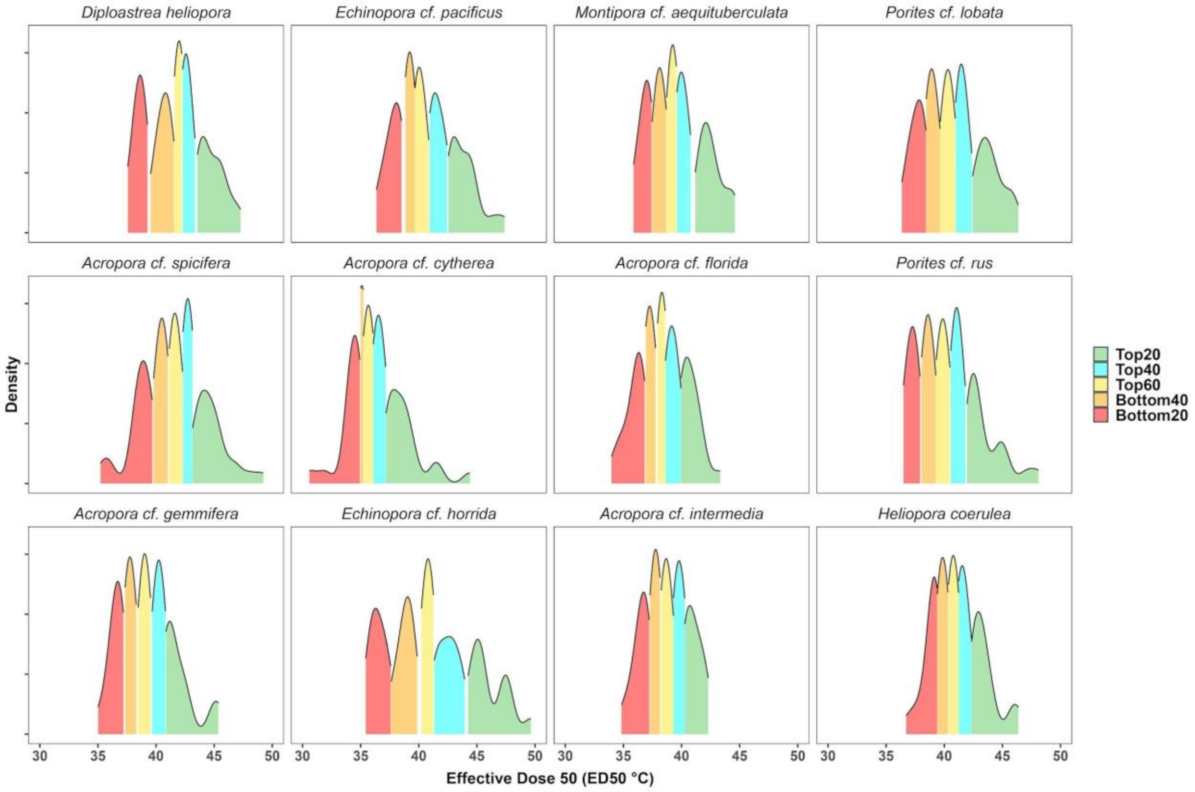


**Supplementary Figure 7. Species-specific density distributions of thermal tolerance threshold temperature (Effective Dose 50, ED 50 in °C) across coral colonies ranked into Top to Bottom quantile groups.** Thermal tolerance declines from top-ranked (green and cyan) to bottom-ranked (orange and red) corals, showing a clear stepwise pattern across all species.

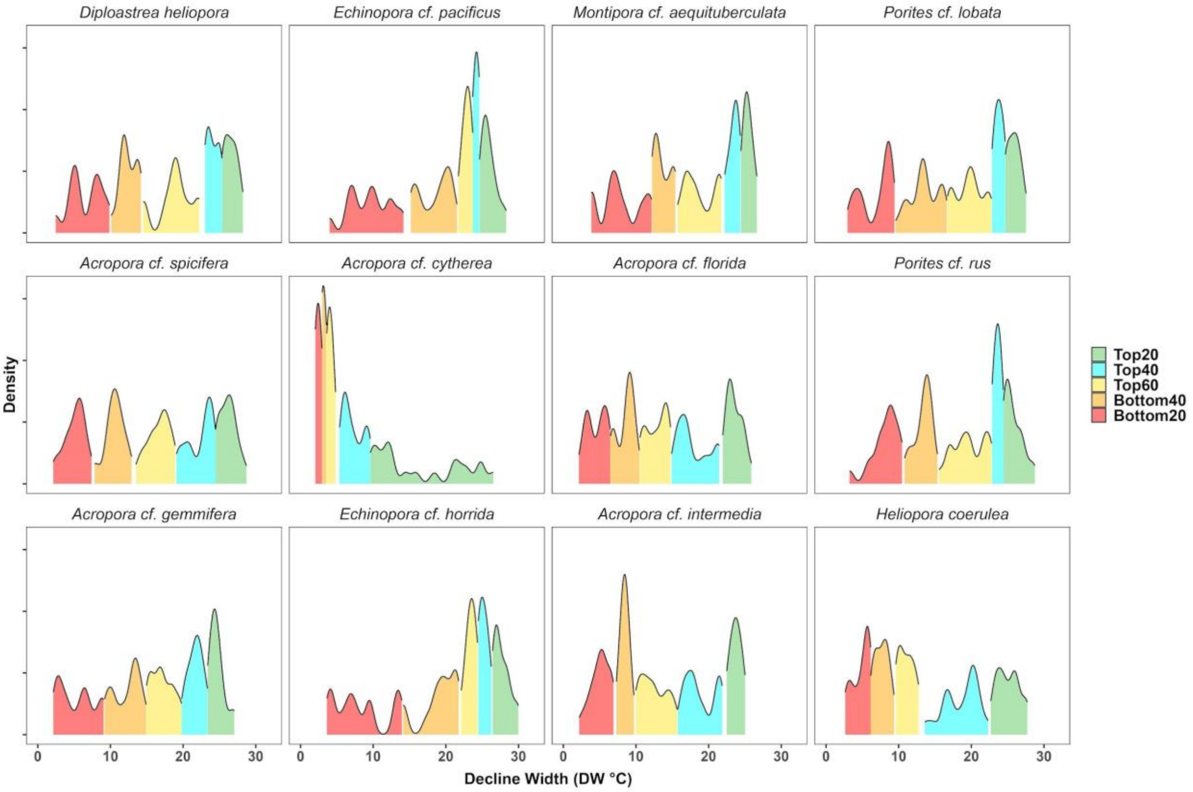


**Supplementary Figure 8. Species-specific density distributions of Decline Width (DW in °C) values across coral colonies ranked into Top to Bottom quantile groups.** The Decline Width is calculated as ED95-ED5 (i.e., thermal limit temperature – thermal breakpoint temperature) and describes whether the effective loss of photosynthetic efficiency is gradually or rapidly lost (i.e., higher values suggestive of higher tolerance). DW declines from top-ranked (green and cyan) to bottom-ranked (orange and red) corals, showing a clear stepwise pattern across all species.

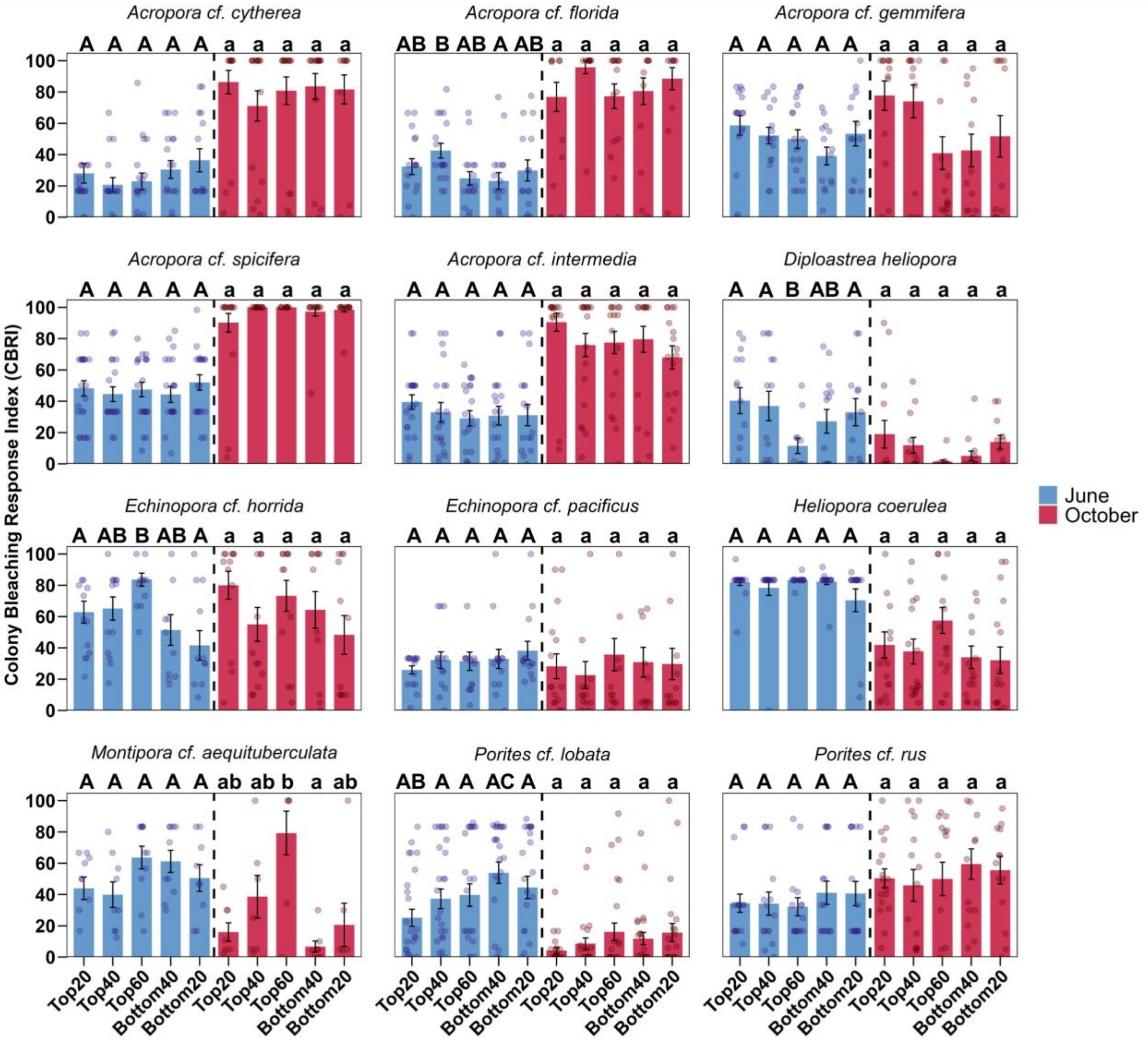


**Supplementary Figure 9. Mean differences and multiple comparisons of coral bleaching severity of coral colonies ranked into top and bottom Effective dose 50 (ED50) quantiles.** Coral bleaching severity (i.e., Colony Bleaching Response Index) was measured in June 2024 during peak heat stress and four months later in October 2024. These measurements reflect immediate heat stress response and subsequent mortality, respectively. ED50 is the temperature threshold at which 50% of the measured photosynthetic efficiency is lost compared to baseline levels. Error bars signify the standard error (±), and dots represent individual data points of coral colonies. Annotated letters alternate in each panel between upper (i.e., June) and lower case (i.e., October) letters for clarity. Letters represent results of pairwise Wilcoxon Rank-Sum Tests conducted at significance levels of $p \leq 0.05$.

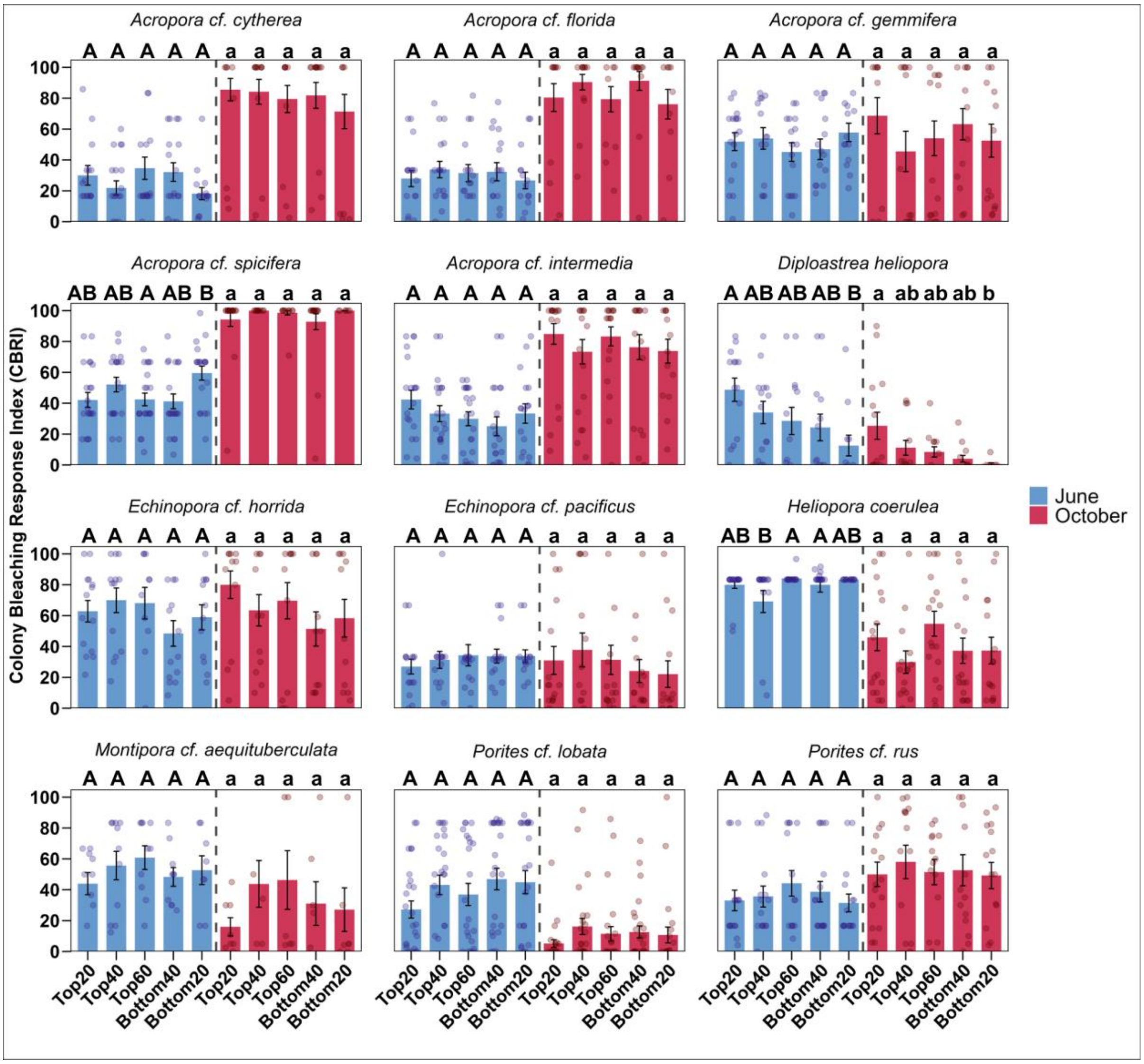


**Supplementary Figure 10. Mean differences and multiple comparisons of coral bleaching severity of coral colonies ranked into top and bottom Decline Width (DW) quantiles.** Coral bleaching severity (i.e., Colony Bleaching Response Index) was measured in June 2024 during peak heat stress and four months later in October 2024. These measurements reflect immediate heat stress response and subsequent mortality, respectively. Decline Width (DW) is calculated as ED95 (Effective Dose 95, temperature limit at which 95% of photosynthetic efficiency is lost) minus ED5 (i.e., the breakpoint temperature at which the decline in photosynthetic efficiency is initiated), whereby higher values are suggestive of a superior ability to cope with heat stress.Error bars signify the standard error (±), and dots represent individual data points of coral colonies. Annotated letters alternate in each panel between upper (i.e., June) and lower case (i.e., October) letters for clarity. Letters represent results of pairwise Wilcoxon Rank-Sum Tests conducted at significance levels of p ≤0.05.

### Tables

**Supplementary Table 1. Overview of the number of surveyed colonies in June (n) and October (N).** Percentage number of colonies observed healthy (i.e., no bleaching), bleached (i.e., >90% of colony surface bleached), and dead (>90% of colony surface dead) are listed for each species during peak heat stress in June 2024 and after heat stress in October 2024.

| | | June 2024 | | | October 2024 | | | |
|---|---|---|---|---|---|---|---|---|
| Species | n | CBRI | Healthy (%) | Fully Bleached (%) | N | CBRI | Healthy (%) | Dead (%) |
| *Acropora* cf. *cytherea* | 167 | 28.6 | 13.2 | 1.8 | 176 | 79.4 | 4.6 | 75.0 |
| *Acropora* cf. *florida* | 177 | 30.9 | 8.5 | 1.7 | 178 | 78.1 | 3.4 | 64.6 |
| *Acropora* cf. *gemmifera* | 160 | 49.0 | 1.3 | 4.4 | 165 | 60.1 | 15.9 | 46.3 |
| *Acropora* cf. *spicifera* | 172 | 45.9 | 4.1 | 6.4 | 173 | 96.8 | 0 | 95.4 |
| *Acropora* cf. *intermedia* | 148 | 33.6 | 7.4 | 8.8 | 151 | 81.4 | 3.3 | 66.2 |
| *Diploastrea heliopora* | 105 | 33.5 | 26.7 | 7.6 | 104 | 10.0 | 55.8 | 0.1 |
| *Echinopora* cf. *horrida* | 158 | 57.5 | 2.5 | 17.1 | 161 | 63.4 | 2.5 | 44.7 |
| *Echinopora* cf. *pacificus* | 173 | 31.7 | 1.2 | 0 | 169 | 30.9 | 11.8 | 13.6 |
| *Heliopora coerluea* | 135 | 80.1 | 1.5 | 86.0 | 134 | 41.6 | 4.5 | 16.4 |
| *Montipora* cf. *aequituberculata* | 120 | 53.3 | 2.5 | 20.8 | 108 | 41.0 | 21.3 | 27.8 |
| *Porites* cf. *lobata* | 197 | 44.8 | 16.8 | 23.4 | 193 | 12.4 | 54.9 | 1.0 |
| *Porites* cf. *rus* | 151 | 33.5 | 2 | 15.2 | 141 | 48.9 | 8.5 | 10.6 |
